# Enhanced efficacy of vaccination with vaccinia virus in old versus young mice

**DOI:** 10.1101/658005

**Authors:** Evgeniya V. Shmeleva, Geoffrey L. Smith, Brian J. Ferguson

**Author notes:** **Correspondence:** Brian J. Ferguson, Geoffrey L. Smith.

## Abstract

Immunosenescence is believed to be responsible for poor vaccine efficacy in the elderly. To overcome this difficulty, research into vaccination strategies and the mechanisms of immune responses to vaccination is required. By analyzing the innate and adaptive immune responses to vaccination with vaccinia virus (VACV) in mice of different age groups, we found that immune cell recruitment, production of cytokines/chemokines and control of viral replication at the site of intradermal vaccination were preserved in aged mice and were comparable with younger groups. Analysis of cervical draining lymph nodes (dLN) collected after vaccination showed that numbers of germinal center B cells and follicular T helper cells were similar across different age groups. The number of VACV-specific CD8 T cells in the spleen and the levels of serum neutralizing antibodies 1 month after vaccination were also comparable across all age groups. However, following intranasal challenge of vaccinated mice, body weight loss was lower and virus was cleared more rapidly in aged mice than in younger animals. In conclusion, vaccination with VACV can induce an effective immune response and stronger protection in elderly animals. Thus, the development of recombinant VACV-based vaccines against different infectious diseases should be considered as a strategy for improving vaccine immunogenicity and efficacy in the elderly.

## 1 Introduction

Old people have increased susceptibility to viral and bacterial infections (1) and in people above 65, about a third of mortality is related to infections (2, 3). Prophylactic vaccination is recommended for the elderly to reduce the burden and severity of infectious diseases (4). However, the elderly respond poorly to the majority of existing vaccines, including vaccines against influenza virus, pneumococcus, hepatitis B, tetanus, pertussis, and diphtheria (5–10). It is important, therefore, to search for ways to overcome this barrier.

The reported decline in the immune system fitness with age, is thought to contribute to reduced vaccine efficacy in humans and mice (5, 6, 11, 12). This decline impacts both innate and adaptive immunity. Impaired recognition of microorganisms and their components, inadequate receptor signaling, and altered cytokine production have all been reported (13). Additionally, dysfunctionality of innate immune cells such as neutrophils, NK cells, monocytes, macrophages and dendritic cells in their ability to migrate, perform phagocytosis, kill bacteria and secrete cytokines have been noted (2, 14–16). Decline in the performance of multiple aspects of the adaptive immune response with age also occurs. This includes decreased numbers of naïve T cells, a reduced TCR repertoire, an impaired clonal expansion and generation of functional effector and memory T cells, a decrease in immunoglobulin class switch recombination, and restricted B cell diversity and antibody production (7, 11, 17).

Vaccinia virus (VACV), a dsDNA poxvirus (18), is the vaccine used to eradicate smallpox (19). VACV replicates in the cytoplasm of infected cells and has a large genome containing approximately 200 genes (20). Between one third and one half of these genes encode proteins dedicated to immune evasion (21). Although VACV is immunosuppressive, vaccination with VACV in humans and mice results in the generation of robust, long-lasting antibody and T-cell memory that provides protection against re-infection (21–25). The ability of VACV to generate such potent humoral and cellular memory, and its proven ability to protect a population against infectious disease, makes it an excellent model system for studying immune response to vaccination. In this study, we use a mouse model of VACV intradermal vaccination that generates protective immunity against re-infection (26). In this model, both antibody and T cell memory responses are robust and consistent and contribute to protection against subsequent challenge with VACV (27, 28). Although VACV has been studied intensively in multiple models, the influence of aging on the immune and vaccination responses to VACV is unexplored. In this study, we analyzed the innate and adaptive immune response to VACV infection and evaluated subsequent resistance to re-infection in three different age groups of mice.

## 2 Materials and Methods

### 2.1 Animals and study design

C57BL/6 female mice were used in the study. All animals were purchased from Charles River and housed in the Cambridge University Biomedical Services facility. All animal experiments were conducted according to the Animals (Scientific Procedures) Act 1986 under PPL 70/8524 issued by UK Home Office.

The animal experiments included intradermal (i.d.) vaccination and intranasal (i.n.) challenge (Fig. 1). Animals of 7, 22 and 54 weeks old (wo) received i.d. injections with 10^4^ plaque-forming unit (PFU) of VACV strain Western Reserve (WR) or control vehicle (0.01% BSA/PBS) into both ear pinnae. VACV used for infection of animals was purified from infected cells by sedimentation through a sucrose cushion and subsequently through a sucrose density gradient. Virus infectious titers were determined by plaque assay on BSC-1 cells and frozen at −70 °C until use. To evaluate the immune response during the acute stage post vaccination, ear tissues and cervical draining lymph nodes (dLN) were collected at day (d) 7 after i.d. infection for measurement of infectious viral titers (by plaque assay), leukocyte infiltration (by FACS) and levels of cytokines/chemokines (by Luminex assay). Serum and spleens were obtained at 29 d post i.d. injections to measure the titers of anti-VACV neutralizing antibodies and the composition of T cell subpopulations.

**Figure 1.**
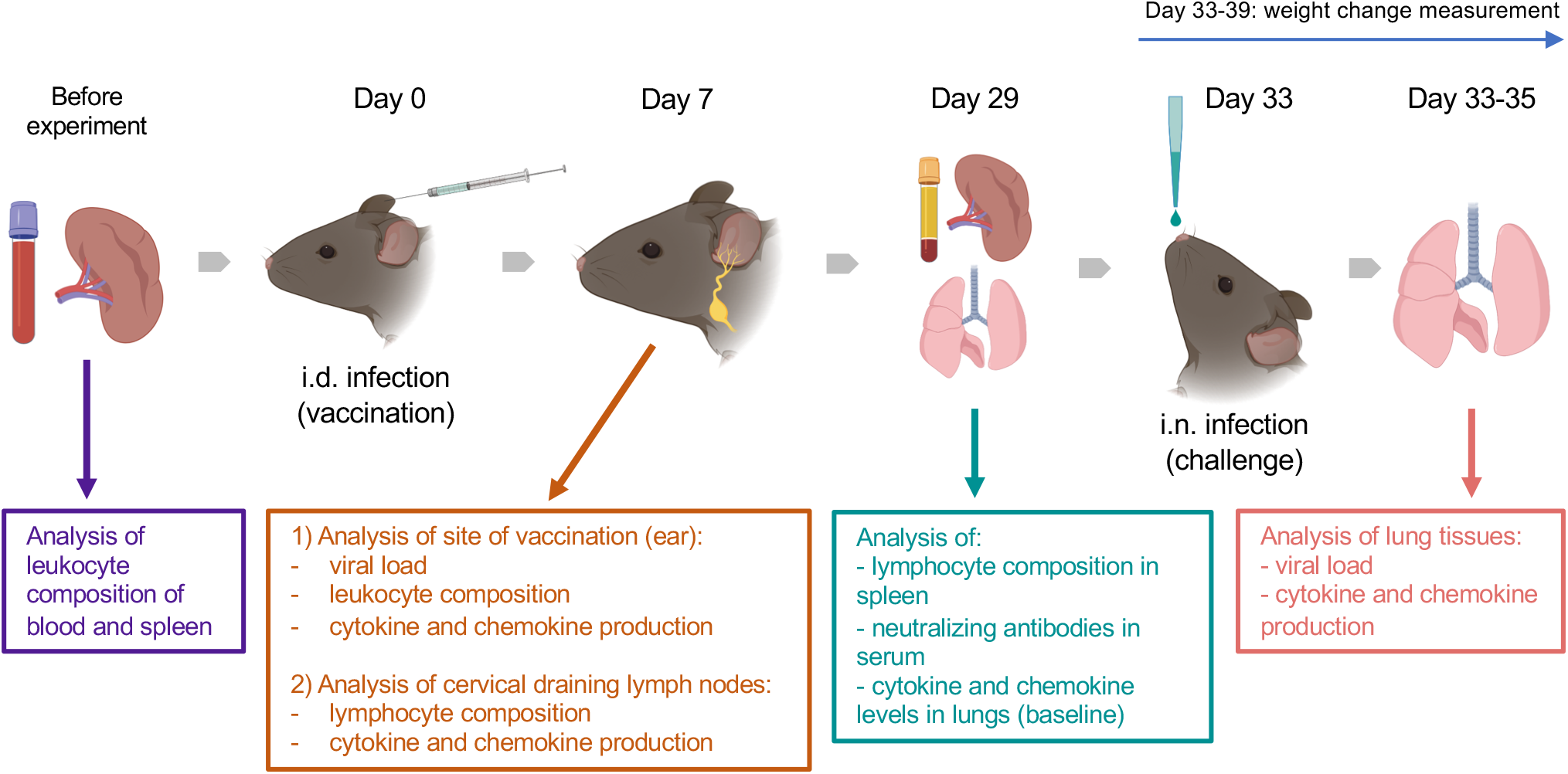
Experimental design. Groups (*n*=4-5) of 7-, 22- and 54-week old C57BL/6 mice were used in the study. Various parameters were measured before and at 7 and 29 d after intradermal (i.d.) infection with 10^4^ PFU of VACV WR, as well as following intranasal (i.n.) challenge of immunised or naïve mice with ~10^7^ PFU of VACV WR.

To assess the efficacy of vaccination, vaccinated mice (33 d post i.d. VACV infection) and naïve (non-vaccinated) mice were challenged i.n. with ~10^7^ PFU of VACV WR. The body weights of animals were monitored daily. Whole lungs were collected at 12, 24 and 48 h post challenge to measure the viral load and the levels of cytokines/chemokines in tissue.

The baseline of immunological parameters was measured in the blood, spleens and lungs of naïve, uninfected animals (n=4).

### 2.2 Flow cytometry

FACS analysis was performed to measure the immune cells present in ear tissue, cervical dLN, blood and spleens of vaccinated and mock-vaccinated animals.

Ear pinnae were collected at 7 d post i.d. infection, then separated into dorsal and ventral layers and both leaflets were placed into 1.5 ml of the RPMI-1640 (Gibco, Cat. # 21875034) medium containing 750 U/ml of collagenase I (Gibco, Life Technologies, Cat. # 17018-029) and 100 U/ml of DNase I (Invitrogen, Cat. # 18047-019), followed by 1 h incubation at 37 °C on an orbital shaker, at 1100 rpm. Suspensions containing digested ear samples were mashed through a 70-μm cell-strainer, mixed with 10 ml of RPMI-1640 medium containing 35% of isotonic Percoll (Sigma, Cat. # P1644-500ML) and centrifugated for 10 min at 940 relative centrifugal force (rcf) without use of brake, at 21 °C. Then the supernatants were removed and the cells were washed with PBS.

To obtain cells from spleen or dLN, organs were mashed through 70-μm cell-strainers and washed with PBS.

Before antibody staining of prepared cell suspensions, red blood cells (RBC) were lysed with BD Pharm Lyse (BD Biosciences, Cat. # 555899) and washed twice. The suspensions were then passed through 70-μm Pre-Separation Filters (Miltenyi, Cat. # 130-095-823) and cells were counted using a NucleoCounter NC-250 (Chemometec).

For the staining of cell surface markers, the samples were incubated with Zombie Fixable Viability dye (Suppl. Table 1) and, after one washing step, purified rat anti-mouse CD16/CD32 antibody (Mouse BD Fc Block) (BD Biosciences, Cat. # 553141) was added to the cell suspension to block non-specific binding. For intracellular Bcl-6 and Ki-67 staining, Foxp3 / Transcription Factor Staining Buffer Set (eBioscience, Cat. # 00-5523-00) was used. Then surface or intracellular markers were stained with monoclonal antibodies (mAbs). The myeloid panel for surface staining of ear tissue included: CD45, Siglec-F, CD11c, CD11b, Ly6C, Ly6G, as well as dump channel markers (CD3, CD5, CD19, NK1.1). The lymphoid cells in ear tissue were identified using mAbs to CD45, NK1.1, CD3, CD4, CD8 and with MHC dextramer H-2Kb/TSYKFESV. For assessment of VACV-specific CD8 T cells in the dLN, the cells were stained with mAbs to CD45, CD19, CD3, CD8 and with MHC dextramer H-2Kb/TSYKFESV. The panel for identification of germinal center B cells and follicular helper T lymphocytes in dLNs included mAbs to CD4, CXCR5, PD-1, B220, Bcl-6 and ki-67. Subpopulations of CD4 and CD8 T cells in spleen were determined by staining with mAbs to CD45, CD3, CD8, CD4, CD62L and CD44 and with MHC dextramer H-2Kb/TSYKFESV. All dyes and mAbs used in the study are listed in Suppl. Table 1. After final washing steps, cells were resuspended with PBS containing 4% paraformaldehyde and were analyzed by FACS on a BD LSRFortessa (BD Biosciences). Gating strategies are shown in Suppl. Figs. 1-5.

For the Trucount assay, blood was collected into Micro K3EDTA Tubes (Sarstedt, Cat. # 41.1395.005) to prevent clot formation. Then, 50 μl of whole blood was pipetted into the bottoms of BD Trucount Tubes (BD Biosciences, Cat. # 340334), followed by 5 min incubation with Mouse BD Fc Block. The samples were then stained with mAbs to CD45, CD3, CD4, CD8, CD19, NK1.1, CD11b and Ly6G (Suppl. Table 1). After RBC lysis, and without washing steps, the absolute numbers of different leukocyte populations were determined by analysis on a BD LSRFortessa. The gating strategy is shown in Suppl. Fig. 6.

### 2.3 Intravascular staining

To discriminate leukocytes resident in ear tissue from cells located in vasculature, intravascular staining was undertaken as described (29). Briefly, 5 mins before culling, mice were given an intravenous infusion into the tail vein of anti-CD45-BV421 mAb (BioLegend, Cat. # 103134). Ears were then collected, and cells were isolated as described under Flow cytometry above. The cell suspension from ear tissue was stained with anti-CD45-PE (BioLegend, Cat. # 103106). Blood leukocytes were gated as double positive (CD45-BV421^+^ CD45-PE^+^) cells, while tissue immune cells were positive only for CD45-PE (see Suppl. Fig. 7).

### 2.4 Identification of cytokines and chemokines in ear, dNL and lung tissues

Whole ears, dLN or lungs were homogenized in 1.5 ml flat-bottom tubes containing 400 μl of 0.5% BSA/PBS using an OMNI Tissue Homogenizer with plastic hard tissue probes (OMNI International). The tissue homogenates were centrifugated at 10,000 rcf for 20 min, at 4 °C and supernatants were obtained and stored at −70 °C. Magnetic Luminex Mouse Premixed Multi-Analyte kits were purchased from R&D Systems, to assess levels of IFNγ, TNFα, IL-1β, IL-4, IL-6, IL-10, IL-33, CCL2, CCL3, CCL4, CCL5, CCL7, CCL20, CXCL1, CXCL2, CXCL5 (LIX) and CXCL10 using a Luminex 200 analyzer (Luminex Corporation).

### 2.5 Measurement of viral loads in ear and lung tissues

Whole ears and lungs were homogenized as described above. The homogenates underwent 3 cycles of freezing-thawing-sonicating to rupture cells and release the virus. Titers of infectious virus in ear samples were then determined by plaque assay using BSC-1 cell monolayers.

The VACV load in the lungs of vaccinated mice was measured by determining the virus genome copy number by qPCR as described (30). Genome copy number correlated well with measurement of virus infectivity by plaque assay (Suppl. Fig. 8). Supernatant samples from lung tissue homogenates were prepared by centrifugation of samples at 1000 rcf for 5 min, followed by 10-fold dilution of supernatants with nuclease-free water (Cat. # AM9930, Ambion). The reaction mix for real-time qPCR included: 2 μl of template, 10 μl of 2x qPCRBIO Probe Mix (Cat. # PB20.21-5, PCRBiosystems), 0.8 μl of 10 μM VACV gene *E9L* forward primer (CGGCTAAGAGTTGCACATCCA), 0.8 μl of 10 μM *E9L* reverse primer (CTCTGCTCCATTTAGTACCGATTCT), 0.4 μl of 10 μM *E9L* probe (TaqMan MGB Probe – AGGACGTAGAATGATCTTGTA, Applied Biosystems). The reaction volume was adjusted to 20 μl with nuclease-free water. A plasmid containing the VACV *E9L* gene served as a standard and was a gift from Brian M Ward, University of Rochester Medical Center, USA. qPCR assays were run on a ViiA 7 Real-Time PCR System (Applied Biosystems) with the following protocol: initial denaturation step at 95 °C for 3 min, followed by 40 cycles of denaturation at 95 °C for 5 sec, annealing and extension at 60 °C for 30 sec.

### 2.6 Assessment of VACV neutralizing anybody titer in serum

Blood was collected into Microvette CB 300 μl tubes with clot activator (Sarstedt, Cat.# 16.440.100). Blood samples were left at room temperature for 2 h to allow clot formation. After centrifugation at 10,000 rcf for 5 min at room temperature, serum was collected and stored at −70 °C. Titers of neutralizing antibodies were assessed by plaque reduction neutralization test as described (31). Serum samples were incubated at 56 °C for 30 min to inactivate complement, then two-fold serial dilutions were prepared (1:50, 1:100, 1:200, 1:400, 1:800 and 1:1600) using 2.5% fetal bovine serum (FBS)/1% PenStrep/DMEM medium. The diluted serum samples, or reference samples (medium only), were mixed 1:1 with medium containing 3.2 × 10^2^ PFU/ml of VACV WR that had been purified by sedimentation through a sucrose density gradient. After 1 h incubation at 37 °C, samples were titrated by plaque assay and half maximal inhibitory concentrations (IC50) were calculated.

### 2.7 Statistical analysis

SPSS v.25 and GraphPad Prism v.8 were used for statistical analysis. The Mann–Whitney U-test was applied for the comparisons of two groups of animals and two-way repeated measures (RM) ANOVA tests were performed for the analysis of time series data. The Spearman’s correlation test was used for relation analysis of variables. P values <0.05 were considered significant.

## 3 Results

Three groups (7, 22 or 54 weeks old [wo]) of female C57BL/6 mice were used representing young adults, middle-aged and old animals. Fifty-four-wo animals were chosen to represent the elderly group based on their general appearance (graying coat, thinning hair) and the death rate in the colony (~10% lethality over a 6-week period not associated with the experiment). One of the features of immunosenescent phenotype is a decline in naïve T cell numbers, which correlates with increased morbidity and mortality (32, 33). Notably, the 54-wo mice had significantly decreased absolute numbers of CD4 and CD8 cells in the blood and spleen in comparison with younger animals (Fig. 2A). This reduction was due to the decline of naïve subpopulations of CD4 and CD8 cells, while effector T cell numbers were increased in 54- and 22-wo mice in comparison with the 7-wo group (Fig. 2B, Suppl. Fig. 5).

**Figure 2.**
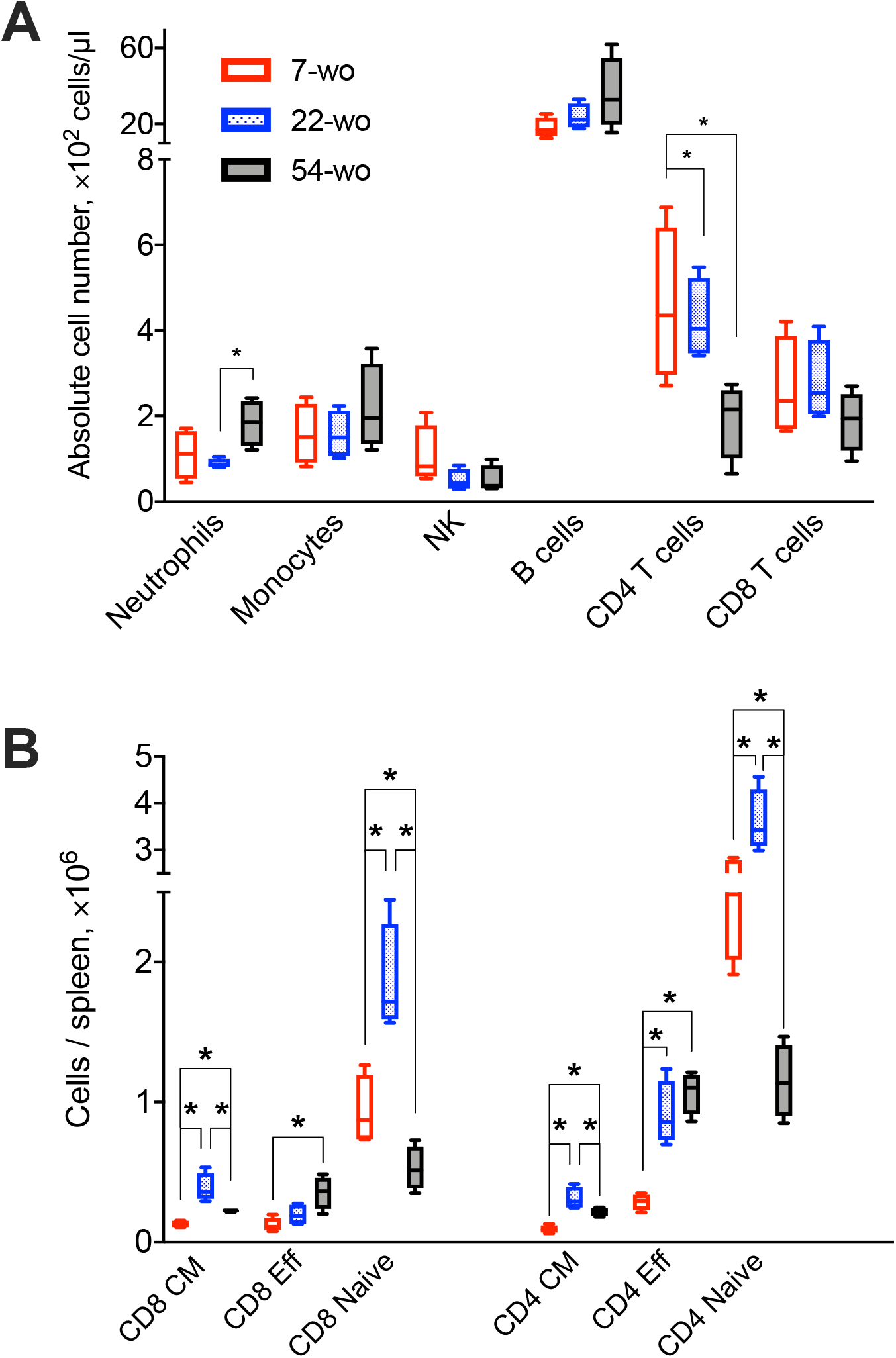
Old mice have decreased numbers of naïve CD8 and CD4 T cells. The absolute numbers of different subpopulation of leukocytes in blood (**A**) and T cells isolated from the spleen (**B**) of 7-, 22- and 54-week old mice (without VACV infection) (4 animals per group). CM, central memory; Eff, effector. P values determined by Mann-Whitney test, * = p<0.05.

### 3.1 Immune response to intradermal infection with VACV is conserved across different age groups

Intradermal (i.d) infection with VACV leads to the development of skin lesions 5-6 days (d) post infection that usually heal within 21 d (26). Using this infection model, immune cell recruitment, the levels of cytokines/chemokines and the viral load in ear tissues 7 d post i.d. infection was analyzed in three different age groups of mice.

FACS analysis of leukocyte populations in the infected ear showed that ~97% represented cells that had infiltrated into the tissue, whereas immune cells from blood circulation constituted only 3% of the total leukocytes (Suppl. Fig. 7). In comparison with 7- and 22-wo mice, infected ear tissues of 54-wo animals showed significantly less infiltration of different leukocyte populations including NK, CD4 and CD8 T cells, Ly6C^+^ (inflammatory) monocytes and CD45^+^CD3^−^CD5^−^CD19^−^NK1.1^−^Siglec-F^−^CD11c^+^ cells, which represent a mixed population of dendritic cells and macrophages (Fig 3A). The presence of VACV-specific CD8 T cells in infected ear tissue of 54-wo mice was also reduced compared to the other groups (Fig. 3B). Given that amounts of CD4 and CD8 T cells in the elderly mice were diminished before the infection (Fig. 2), the lower numbers of lymphoid populations infiltrating ears are likely reflective of the reduced availability of T cells in the blood circulation.

**Figure 3.**
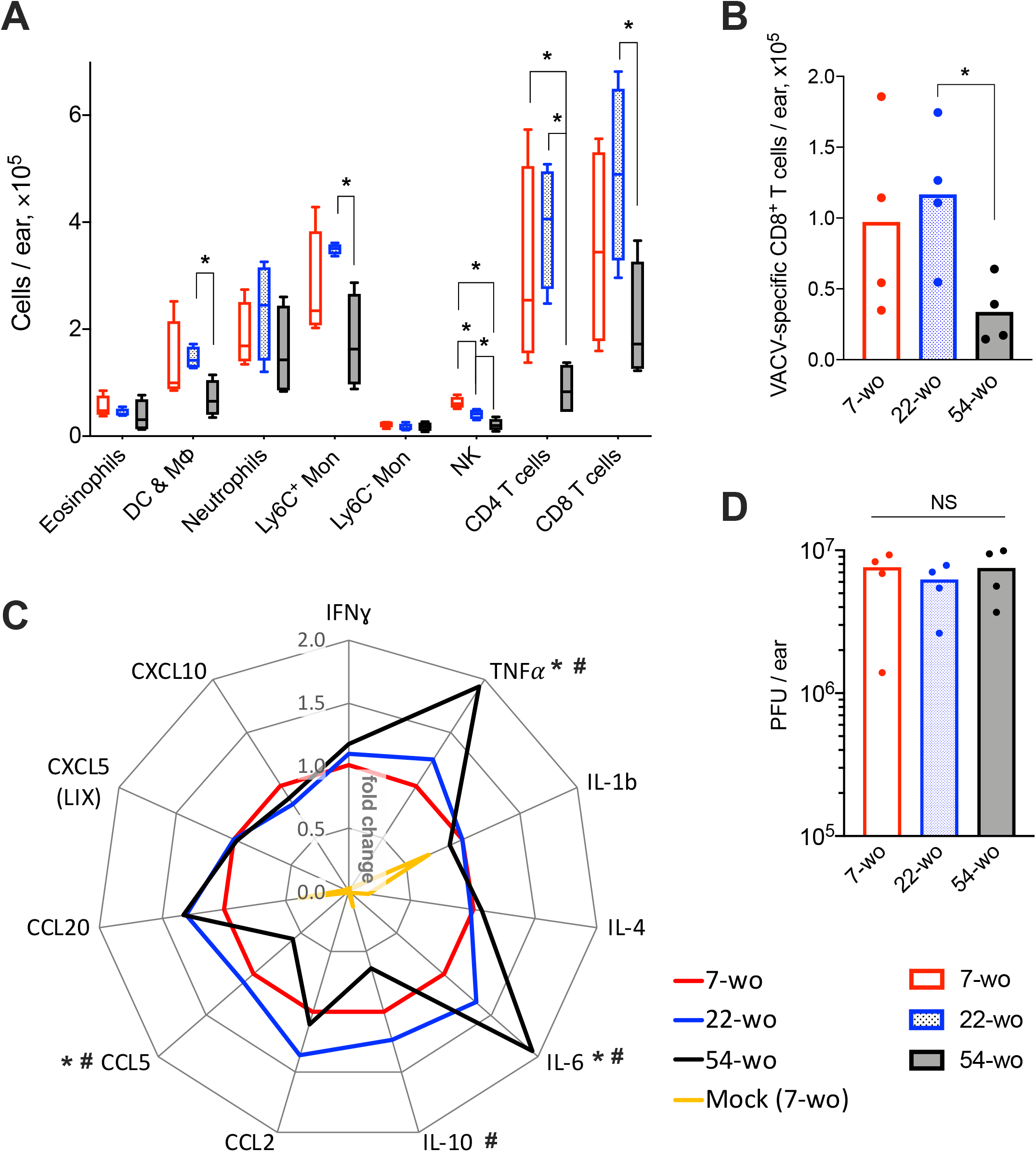
Conservation of the immune response to intradermal infection with VACV across age groups. Ear tissues were collected at 7 d post i.d. infection with VACV or PBS (mock-control) from groups of 7-, 22- and 54-wo mice (*n*=4-5 per group). The absolute numbers of (**A**) different subpopulations of leukocytes and (**B**) VACV-specific CD8 T cells infiltrating ear tissues are shown. DC & MФ, dendritic cells and macrophages; Mon, monocytes; p values were determined by the Mann-Whitney test, * = p<0.05. (**C**) The levels of cytokines/chemokines detected by multiplex assay (Luminex) in ear tissues are presented as fold change compared with the 7-wo VACV-infected group, which is assigned a value of 1. The means are shown; p values were determined by the Mann-Whitney test, * = p<0.05 between 7- and 54-wo animals. # = p<0.05 between 22- and 54-wo mice. (**D**) Titers of VACV in ear tissues at 7 d post i.d. infection with VACV. PFU, plaque-forming units; NS – non-significant by Mann-Whitney test. The experiment was performed twice and representative data from one experiment are shown.

The levels of cytokines and chemokines detected in ear tissue of the old animals at 7 d after i.d. infection did not differ greatly from the young and middle-age groups (Fig. 3C). Only the levels of IL-10 and CCL5 were reduced, while the concentrations of TNFα and IL-6 were increased in 54-wo mice in comparison with 7- and 22-wo animals. However, the amplitude of these changes did not exceed 2-fold. In addition, viral loads in infected ear tissues were similar in all groups (Fig. 3D). Hence, the infected ear tissue was able to respond to the infection via production of inflammatory mediators and control virus infection to equivalent levels across all age groups. These data provide further evidence that the lower cell recruitment into the ear tissue of 54-wo mice was likely due to the reduced availability of circulating cells rather than due to changes in local responses in the infected tissue. Thus, functionally, the immune response to i.d. vaccination with VACV was preserved in 54-wo mice, and the ability to control VACV replication at the site of infection was equal across 7-, 22- and 54-wo groups.

### 3.2 54-week old mice have enhanced cytokine response to intradermal VACV infection in draining lymph nodes

To investigate the effect of age on the adaptive immune response to vaccination with VACV, cervical draining lymph nodes (dLN) were analyzed at 7 d post i.d. infection. This showed a trend in reduction of absolute numbers of VACV-specific CD8 T cells in the 54-wo mice in comparison with 22- and 7-wo groups (Fig. 4A). However, the amounts of germinal center (GC) B cells and T follicular helper (Tfh) lymphocytes were not significantly different between all groups (Fig. 4B, C).

**Figure 4.**
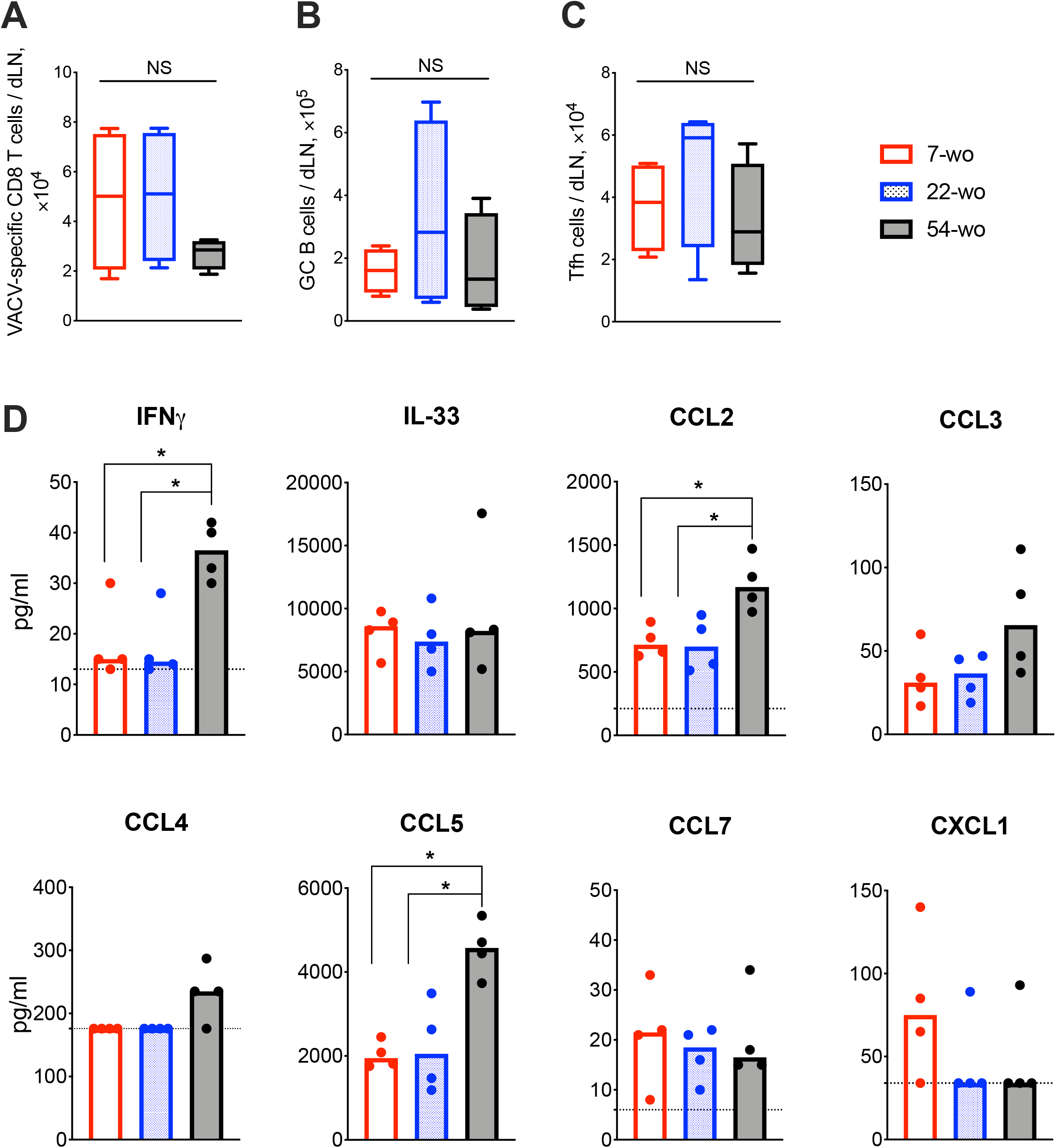
Enhanced cytokine production in the draining lymph nodes (dLN) of 54-wo mice following intradermal infection with VACV. Groups of 7-, 22- and 54-wo mice (*n*=4-5 per group) were infected i.d. with VACV and at 7 d post infection the dLN were collected. The absolute number of (**A**) VACV-specific CD8 T cells, (**B**) germinal center B cells and (**C**) T follicular helper cells were determined by FACS. GC, germinal center; Tfh, T follicular helper; NS, non-significant by Mann-Whitney test. (**D**) The levels of cytokines and chemokines were detected by multiplex assay (Luminex) in cervical dLN from mice treated as above. Medians are shown; dashed lines indicate limit of sensitivity; p values were determined by the Mann-Whitney test, * = p<0.05. The experiment was performed twice and representative data from one experiment are shown.

Next, the levels of cytokines and chemokines in dLN were measured at 7 d post vaccination. Amongst 17 different molecules assessed, IFNγ, IL-33, CCL2, CCL3, CCL4, CCL5, CCL7 and CXCL1 were detectable (Fig. 4D) and the levels of IFNγ, CCL2 and CCL5 were significantly higher in 54-wo animals than in younger mice, while the others were similar in all groups. Thus, the dLNs of old animals responded well to VACV vaccination, expressing high levels of inflammatory mediators and generating appropriate cellular adaptive immune responses.

### 3.3 VACV vaccination induces strong adaptive immune response in mice of different ages

Next, we compared the cellular and humoral memory responses induced by vaccination with VACV. Spleens and blood samples were obtained from mice 29 d post vaccination of 7-, 22- and 54-wo as well as from mock-vaccinated animals. The total numbers of splenic CD8 T cells were equivalent within the three vaccinated groups, while the absolute numbers of CD4 T cells were slightly reduced in 54-wo mice in comparison with in 7- and 22-wo groups (Fig. 5A). The most pronounced changes in numbers of splenic CD8 and CD4 T subsets were observed for effector T cells. In comparison with baseline parameters before vaccination, effector CD8 and CD4 T lymphocytes increased considerably as a result of vaccination for all groups of mice (Fig. 2B, Fig. 5B). Notably, effector T cells in the 54-wo group expanded proportionally greater than in younger groups. However, analysis of VACV-specific CD8 T cells showed that their absolute counts were comparable within all age groups (Fig. 5C). As for the humoral immune response, the ability of serum to neutralize VACV was identical in all three groups (Fig. 5D). These observations show that vaccination with VACV generates memory immunity in 54-wo mice that is quantitively indistinguishable from that generated in 7- or 22-wo mice.

**Figure 5.**
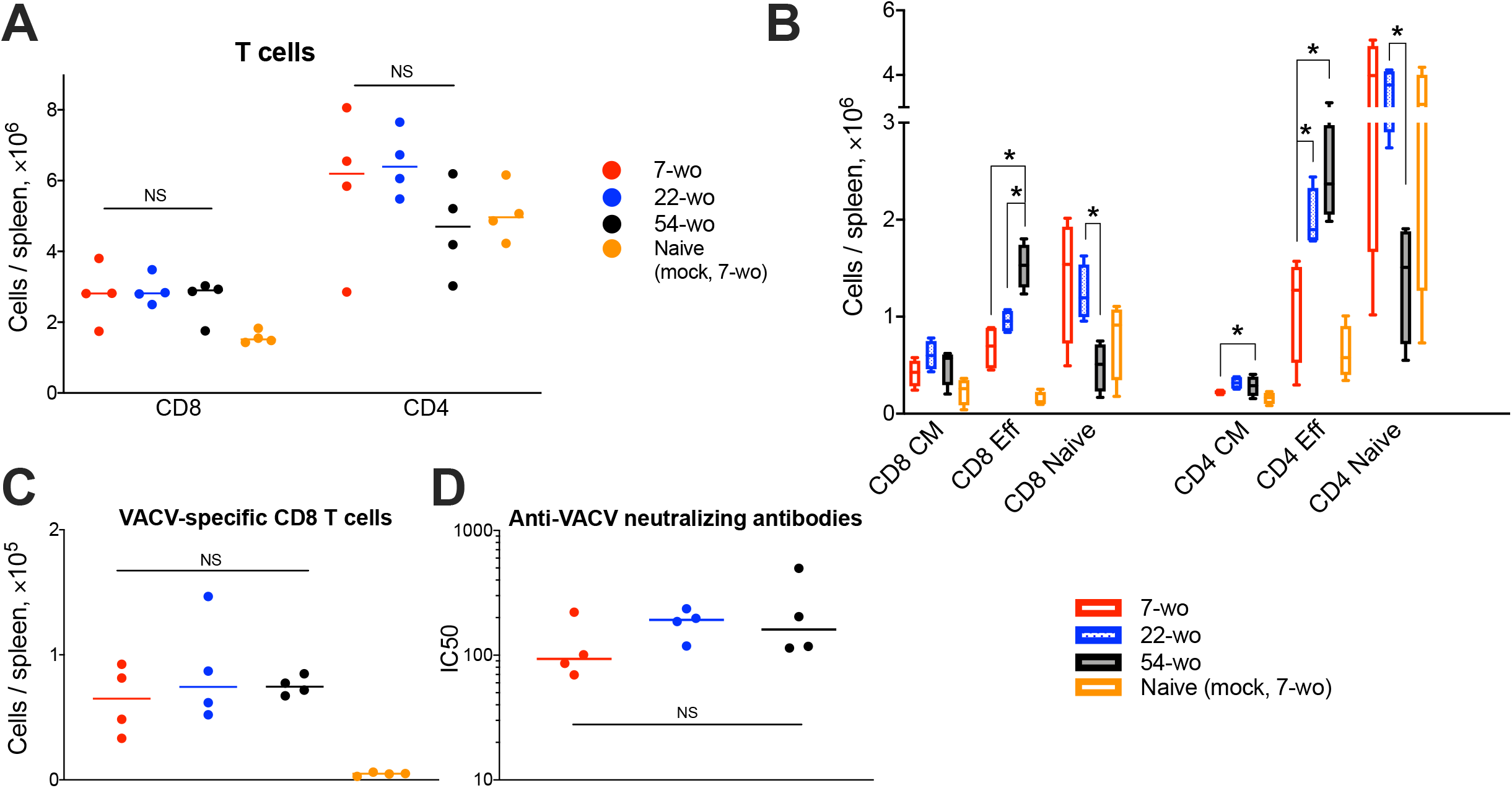
Vaccination with VACV induces a robust adaptive immune response in mice of different ages. Spleens and serum samples were obtained from vaccinated and naïve (mock-vaccinated) mice of different ages at 29 d post i.d. injection (*n*=4-5 animals per group). (**A**) The absolute numbers of total splenic CD8 and CD4 T cells, and (**B**) their subpopulations are shown. Naive, central memory (CM) and effector (Eff). (**C**) Shows VACV-specific CD8 T cells. p values were determined by the Mann-Whitney test, * = p<0.05. NS, non-significant. (**D**) Neutralizing antibody responses determined by plaque-reduction neutralization test. IC50, half maximal inhibitory concentration; NS, non-significant by Mann-Whitney test. All experiments were performed twice and representative data from one experiment are shown.

### 3.4 54-week old mice are better protected against VACV intranasal challenge than those from younger groups

To measure the ability of vaccinated groups to respond to re-infection, the three age groups of vaccinated animals and young naïve mice were challenged i.n. with a dose of VACV equivalent to ~ 300 LD50. All vaccinated groups had mild or moderate weight loss (about 15% maximum) after challenge followed by full recovery. In contrast, naïve mock-vaccinated mice had >25% weight loss and were culled at humane endpoint (Fig. 6A). Notably, following challenge, the 54-wo mice lost less body weight and recovered faster than young and middle-aged groups. Results of viral load measurement in the lungs of challenged mice indicated that the 54-wo mice cleared the virus faster than other groups (Fig. 6B). For the majority of immunized elderly mice, no VACV genome copies were detected in lungs at 24 h post i.n. challenge. Interestingly, the 22-wo mice were slower than 7- and 54-wo groups at clearing the virus, despite the weight loss post challenge being similar between the 7- and 22-wo groups (Fig. 6A, B).

**Figure 6.**
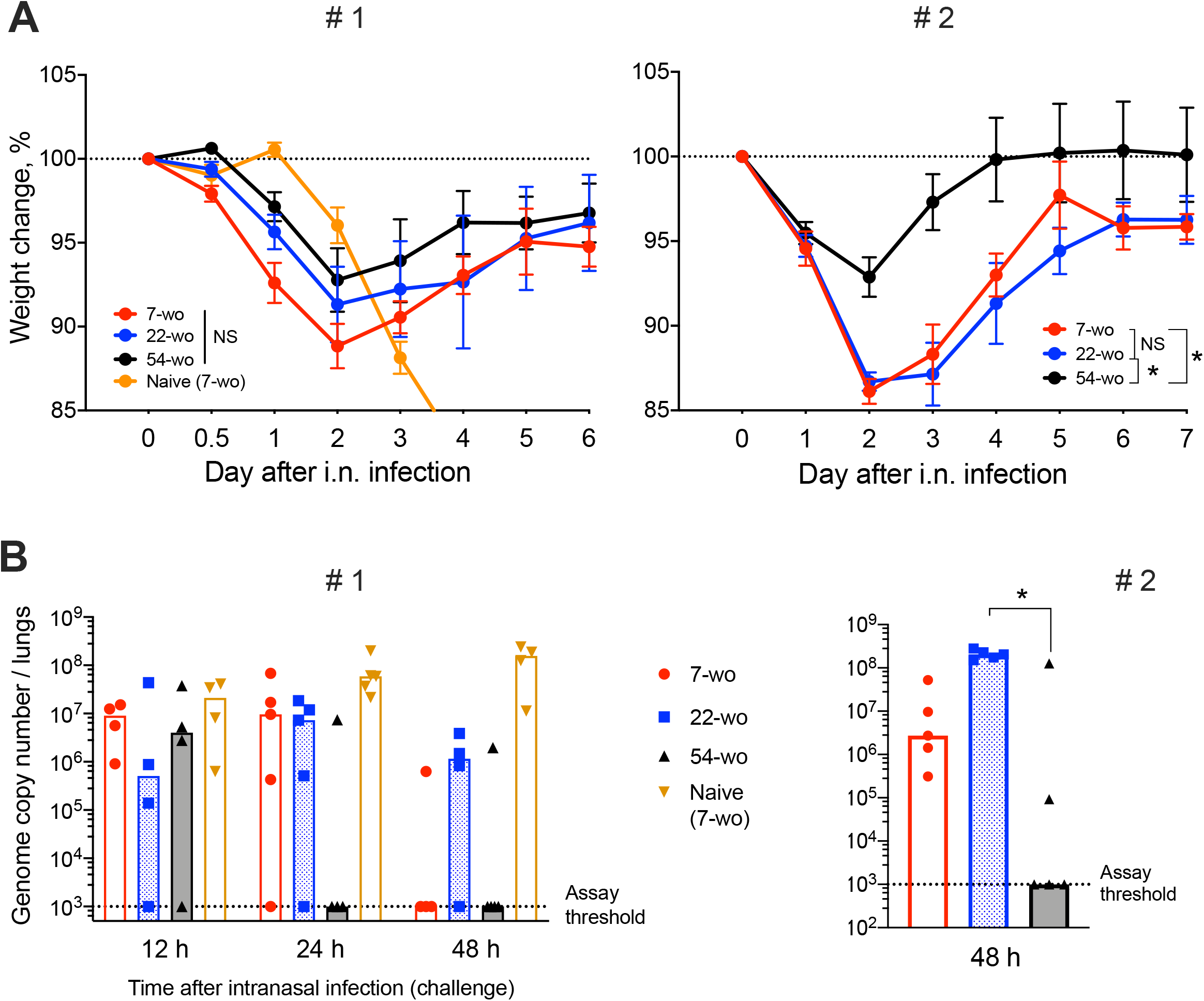
54-week old mice are better protected against VACV intranasal challenge than mice from younger groups. Groups of 7-, 22- and 54-wo C57BL/6 mice (*n*=4-5 per group) were injected intradermal with 10^4^ PFU (per ear, into both ears) of VACV or PBS (mock). These groups were then challenged i.n. with 0.7 × 10^7^ PFU of VACV WR at day 33 post vaccination in experiment #1 (left) and with 1.3 × 10^7^ PFU of VACV WR in experiment #2 (right). **(A)** Body weight changes of mice after intranasal challenge with VACV; within each group, data show a comparison of the weight or each mouse with the weight of the same animal on day zero. The percentages for each group are means with SEM. Statistical analysis by RM ANOVA test. NS, non-significant; * = p<0.05. (**B**) Viral genome copy number in both lungs from mice at 12, 24 and 48 h post i.n. challenge were determined by qPCR. Medians are shown.

Lastly, the levels of inflammatory mediators in the lung tissue of vaccinated and naïve mice were measured (Fig. 7, Suppl. Fig. 9). Baseline levels before i.n. infection did not vary significantly between groups. In comparison with the naïve mice, all vaccinated animals responded very quickly to i.n infection. At just 12 h post-challenge, the levels of IFNγ, CCL7, CXCL1, CXCL2, CXCL10 rose substantially (Suppl. Fig. 9), although there was little variation between the different age groups. Only CXCL1 was increased in the elderly mice, and IFNγ levels tended to be higher in the old and middle-aged mice than in young animals. At 24 h post infection, the majority of lung cytokines and chemokines were reduced in the elderly mice compared with other vaccinated age groups. Moreover, at 48 h after challenge, the levels of inflammatory mediators in the 54-wo group were decreased further and started returning to their initial (baseline) concentrations. This may reflect the faster virus clearance. Therefore, the results of weight loss measurement, viral load and cytokine dynamics in lungs indicate that mice vaccinated at the age of 54-wo had robust protection against reinfection with a lethal dose of VACV, and this protection was even stronger than in the mice vaccinated when 7- and 22-wo.

**Figure 7.**
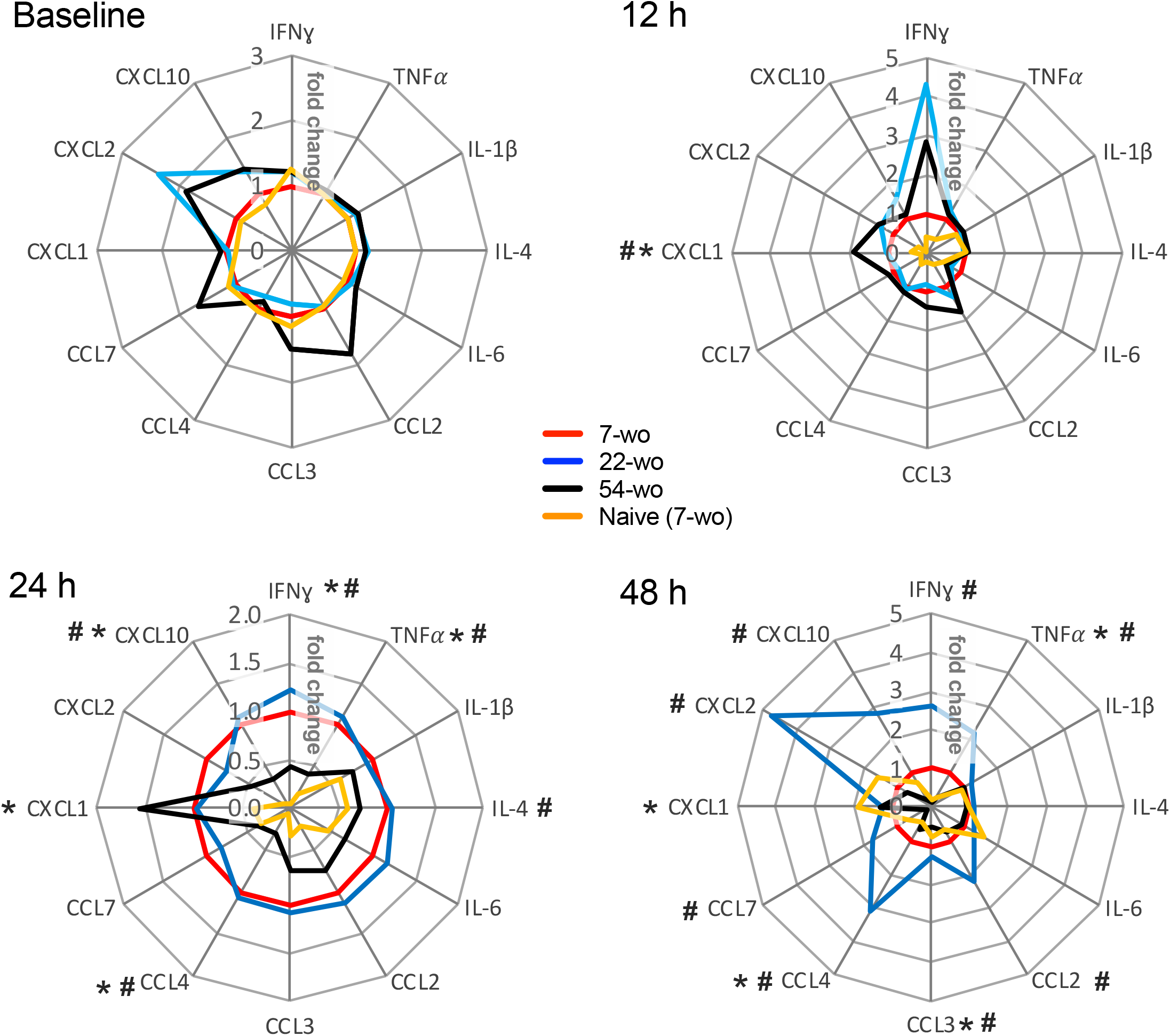
Kinetics of cytokine and chemokine production in lungs after intranasal challenge with VACV correlates with the kinetics of virus clearance. Groups (*n*=4-5) of 7-, 22- and 54-wo C57BL/6 mice were vaccinated i.d. with 10^4^ PFU (per ear, into both ears) of VACV or PBS (mock-control). Then 33 d post vaccination animals were challenged i.n. with VACV WR. Lungs of vaccinated and mock-vaccinated mice were collected at 12, 24 and 48 h after challenge. The levels of cytokines and chemokines were measured by multiplex assay (Luminex). Data are shown as the fold change from the vaccinated 7-wo group, which is assigned a value of 1. Means are shown; p values were determined by the Mann-Whitney test. * = p<0.05 between 7- and 54-wo animals, # = p<0.05 between 22- and 54-wo mice. The experiment was performed twice and data from one representative experiment are shown.

## 4 Discussion

In the current study, we show that i.d. vaccination with VACV leads to successful development of immunological memory in old mice bearing an immunosenescent phenotype. Surprisingly, despite a general reduction of naïve CD8 and CD4 cells (Fig. 2), reduced recruitment of immune cells into the site of i.d. infection (Fig. 3A, B) and inflammatory signatures characteristic of a phenomenon sometimes known as inflammaging (Fig. 3C and 4D), 54-week-old mice demonstrated better vaccination efficacy against challenge than the younger animals.

Vaccination of humans with VACV results in long lasting immunological memory even after administration of a single dose of vaccine (23, 24, 34), and its high efficacy has been validated by the eradication of smallpox. Little information is available concerning VACV vaccine performance in elderly people or mice. One study has reported that vaccination of aged BALB/c mice with recombinant VACV expressing influenza hemagglutinin was effective in generating anti-hemagglutinin antibodies and influenza-specific cytotoxic T cells (35). The basis of high immunogenicity of VACV is not known. However, local immunosuppression by VACV allows the virus to replicate at the site of infection for at least 12 d post i.d vaccination (26). This extended replication period provides constant antigen exposure to the immune system, probably facilitating the generation of strong immunological memory. This immune suppression may be mediated by the scores of immune modulatory proteins expressed by VACV early after infection (21, 36). Many VACV immunomodulators target pattern recognition receptor and interferon receptor signaling to block anti-viral responses in infected cells. In vaccination models, deletion of two or three such genes leads to enhanced safety but decreased immunogenicity of vaccine and impaired protection against challenge (28). The highly attenuated VACV strain modified vaccinia Ankara (MVA), which is replication deficient in many cell types, results in the generation of significantly lower antibody titers in comparison with WR (37, 38). This may explain the potency of VACV in developing robust immunological memory even in old mice. However, this does not explain why better protection against challenge was observed in older mice than in younger counterparts.

One of the features of the immune system in the elderly is the presence of chronic, low grade inflammation, which is sometimes called inflammaging (39). The characteristics of this phenomenon are upregulated activity of NF-κB (40, 41), increased levels of proinflammatory cytokines and chemokines such as TNFα, IL-1, IL-6, IL-8, IL-12, CCL2, CXCL10 (42), accumulation of damage-associated molecular patterns and dysfunctional organelles (43), as well as changes in gut microbiota and metabolism (44). This phenomenon may go some way in explaining the enhanced production of NF-κB-regulated cytokines TNFα and IL-6 in the VACV-infected ear tissue of 54-wo mice (Fig. 3C) as well as CCL2 in dLNs (Fig. 4D). IL-6 and TNF superfamily ligands act as adjuvants and increase immunogenicity of vaccines (45, 46). Therefore, in the case of VACV vaccination, inflammaging might be beneficial by providing additional pro-inflammatory stimulus to drive the cascade of events leading to immune memory development.

Increased production of TNFα and IL-6 along with low levels of IL-10 (Fig. 3C) in old animals after i.d. vaccination, might compensate for the reduced recruitment of immune cells into the vaccination site (Fig. 3A). This might contribute to the control of virus infection after i.d. infection (Fig. 3D) and provide adequate conditions for the generation of immunological memory. Notably, the numbers of Tfh and GC B cells in dLN (Fig. 4B, C), as well as VACV-specific CD8 T cells in the spleen and neutralizing antibody levels (Fig. 5C, D), were similar across all age groups. Nonetheless, it is unclear how the old mice achieved faster clearance of VACV and reduced weight loss after challenge (Fig. 6A, B).

Our results show that absolute counts of splenic effector CD8 T cells expanded substantially and proportionally higher in the elderly group than in the younger mice (Fig. 2B and 5B). This cannot be explained simply by the increased numbers of splenic VACV-specific CD8 T cells (that have been identified by MHC-I dextramer staining) because their absolute counts are too low and similar across all three age groups (Fig. 5C). This difference might be due to the expansion of VACV-specific CD8 T cells against different VACV epitopes and/or the expansion of low-affinity CD8 T cells, which have not been recognized by the type of MHC-I dextramers used in this study. These cells may contribute to the rapid clearance of VACV in the elderly after challenge. Also, despite numerous publications describing functional inefficiency of senescent effector T cells, there are reports that effector T cells from elderly people can have superior immune response to antigen stimulation than younger counterparts (47–49).

In conclusion, this study demonstrates that vaccination of elderly mice is very efficient and not inferior to younger animals. Immunescenescence and inflammaging may be more accurately viewed as immunoadaptation and immunoremodeling in old age, rather than just a slow decline in immune system function (50, 51). The majority of vaccines were created for, and are used in, children and young adults (6), and vaccines designed for the elderly population are needed that consider the specific characteristics of immune system in old age. Given the performance of VACV vaccination shown in the current study, further investigation to understand the mechanisms of its high immunogenicity is warranted.

## 5 Ethics Statement

This study was carried out in accordance with the regulations of The Animals (Scientific Procedures) Act 1986. All protocols and procedures were approved by the UK Home Office and performed under the project licence PPL 70/8524.

## 7 Author Contributions

GS and BF provided the funding. ES, GS and BF designed the study. ES performed all experiments and statistical analysis. GS and BF supervised the work. ES, BF and GS wrote the manuscript.

## 9 Acknowledgments

We thank Dr Brian M Ward (University of Rochester Medical Center, USA) for providing a plasmid containing the VACV *E9L* gene. Also, we thank Dr Michelle A Linterman from the Babraham Institute, Cambridge, UK for helpful discussion.

## 10 Data Availability Statement

Raw data supporting the conclusions of this manuscript will be made available by the authors, without undue reservation, to any qualified researcher.

## References

1. Hepper, H. J., Sieber, C., Cornel, S., Walger, P., Peter, W., Bahrmann, P. et al. Infections in the elderly. Crit Care Clin (2013) 29(3):757–774. doi: 10.1016/j.ccc.2013.03.016.

2. Haq, K., and McElhaney, J. E. Ageing and respiratory infections: the airway of ageing. Immunol Lett (2014) 162(1 Pt B):323–328. doi: 10.1016/j.imlet.2014.06.009.

3. Kline, K. A., and Bowdish, D. M. Infection in an aging population. Curr Opin Microbiol (2016) 2963–67. doi: 10.1016/j.mib.2015.11.003.

4. Weinberger, B. Vaccines for the elderly: current use and future challenges. Immun Ageing (2018) 153. doi: 10.1186/s12979-017-0107-2.

5. Chen, W. H., Kozlovsky, B. F., Effros, R. B., Grubeck-Loebenstein, B., Edelman, R., and Sztein, M. B. Vaccination in the elderly: an immunological perspective. Trends Immunol (2009) 30(7):351–359. doi: 10.1016/j.it.2009.05.002.

6. Ciabattini, A., Nardini, C., Santoro, F., Garagnani, P., Franceschi, C., and Medaglini, D. Vaccination in the elderly: The challenge of immune changes with aging. Semin Immunol (2018) 4083–94. doi: 10.1016/j.smim.2018.10.010.

7. Goodwin, K., Viboud, C., and Simonsen, L. Antibody response to influenza vaccination in the elderly: a quantitative review. Vaccine (2006) 24(8):1159–1169. doi: 10.1016/j.vaccine.2005.08.105.

8. Tin Tin Htar, M., Stuurman, A. L., Ferreira, G., Alicino, C., Bollaerts, K., Paganino, C. et al. Effectiveness of pneumococcal vaccines in preventing pneumonia in adults, a systematic review and meta-analyses of observational studies. PLoS One (2017) 12(5):e0177985. doi: 10.1371/journal.pone.0177985.

9. Weston, W. M., Friedland, L. R., Wu, X., and Howe, B. Vaccination of adults 65 years of age and older with tetanus toxoid, reduced diphtheria toxoid and acellular pertussis vaccine (Boostrix(®)): results of two randomized trials. Vaccine (2012) 30(9):1721–1728. doi: 10.1016/j.vaccine.2011.12.055.

10. Yang, S., Tian, G., Cui, Y., Ding, C., Deng, M., Yu, C. et al. Factors influencing immunologic response to hepatitis B vaccine in adults. Sci Rep (2016) 627251. doi: 10.1038/srep27251.

11. Goronzy, J. J., and Weyand, C. M. Understanding immunosenescence to improve responses to vaccines. Nat Immunol (2013) 14(5):428–436. doi: 10.1038/ni.2588.

12. Solana, R., Pawelec, G., and Tarazona, R. Aging and innate immunity. Immunity (2006) 24(5):491–494. doi: 10.1016/j.immuni.2006.05.003.

13. Molony, R. D., Malawista, A., and Montgomery, R. R. Reduced dynamic range of antiviral innate immune responses in aging. Exp Gerontol (2018) 107130–135. doi: 10.1016/j.exger.2017.08.019.

14. Gomez, C. R., Nomellini, V., Faunce, D. E., and Kovacs, E. J. Innate immunity and aging. Exp Gerontol (2008) 43(8):718–728. doi: 10.1016/j.exger.2008.05.016.

15. Metcalf, T. U., Cubas, R. A., Ghneim, K., Cartwright, M. J., Grevenynghe, J. V., Richner, J. M. et al. Global analyses revealed age-related alterations in innate immune responses after stimulation of pathogen recognition receptors. Aging Cell (2015) 14(3):421–432. doi: 10.1111/acel.12320.

16. Shaw, A. C., Goldstein, D. R., and Montgomery, R. R. Age-dependent dysregulation of innate immunity. Nat Rev Immunol (2013) 13(12):875–887. doi: 10.1038/nri3547.

17. Henry, C., Zheng, N. Y., Huang, M., Cabanov, A., Rojas, K. T., Kaur, K. et al. Influenza Virus Vaccination Elicits Poorly Adapted B Cell Responses in Elderly Individuals. Cell Host Microbe (2019) 25(3):357–366.e6. doi: 10.1016/j.chom.2019.01.002.

18. Moss, B. (2013), Poxviridae, in Fields Virology, edited by B. N. Fields, D. M. Knipe, and P. M. Howley, pp. 2129–2159, Philadelphia: Wolters Kluwer Health/Lippincott Williams & Wilkins,

19. Fenner, F., D. A. Henderson, I. Arita, Z. Jezek, and I. D. Ladnyi (1988), Smallpox and its eradication, 1460 pp., World Health Organization,

20. Goebel, S. J., Johnson, G. P., Perkus, M. E., Davis, S. W., Winslow, J. P., and Paoletti, E. The complete DNA sequence of vaccinia virus. Virology (1990) 179(1):247–66, 517.

21. Smith, G. L., Benfield, C. T., Maluquer de Motes, C., Mazzon, M., Ember, S. W., Ferguson, B. J. et al. Vaccinia virus immune evasion: mechanisms, virulence and immunogenicity. J Gen Virol (2013) 94(Pt 11):2367–2392. doi: 10.1099/vir.0.055921-0.

22. Belyakov, I. M., Earl, P., Dzutsev, A., Kuznetsov, V. A., Lemon, M., Wyatt, L. S. et al. Shared modes of protection against poxvirus infection by attenuated and conventional smallpox vaccine viruses. Proc Natl Acad Sci U S A (2003) 100(16):9458–9463. doi: 10.1073/pnas.1233578100.

23. Crotty, S., Felgner, P., Davies, H., Glidewell, J., Villarreal, L., and Ahmed, R. Cutting edge: long-term B cell memory in humans after smallpox vaccination. J Immunol (2003) 171(10):4969–4973.

24. Taub, D. D., Ershler, W. B., Janowski, M., Artz, A., Key, M. L., McKelvey, J. et al. Immunity from smallpox vaccine persists for decades: a longitudinal study. Am J Med (2008) 121(12):1058–1064. doi: 10.1016/j.amjmed.2008.08.019.

25. Xu, R., Johnson, A. J., Liggitt, D., and Bevan, M. J. Cellular and humoral immunity against vaccinia virus infection of mice. J Immunol (2004) 172(10):6265–6271.

26. Tscharke, D. C., and Smith, G. L. A model for vaccinia virus pathogenesis and immunity based on intradermal injection of mouse ear pinnae. J Gen Virol (1999) 80(Pt 10):2751–2755. doi: 10.1099/0022-1317-80-10-2751.

27. Sumner, R. P., Ren, H., and Smith, G. L. Deletion of immunomodulator C6 from vaccinia virus strain Western Reserve enhances virus immunogenicity and vaccine efficacy. J Gen Virol (2013) 94(Pt 5):1121–1126. doi: 10.1099/vir.0.049700-0.

28. Sumner, R. P., Ren, H., Ferguson, B. J., and Smith, G. L. Increased attenuation but decreased immunogenicity by deletion of multiple vaccinia virus immunomodulators. Vaccine (2016) 34(40):4827–4834. doi: 10.1016/j.vaccine.2016.08.002.

29. Anderson, K. G., Mayer-Barber, K., Sung, H., Beura, L., James, B. R., Taylor, J. J. et al. Intravascular staining for discrimination of vascular and tissue leukocytes. Nat Protoc (2014) 9(1):209–222. doi: 10.1038/nprot.2014.005.

30. Baker, J. L., and Ward, B. M. Development and comparison of a quantitative TaqMan-MGB real-time PCR assay to three other methods of quantifying vaccinia virions. J Virol Methods (2014) 196126–132. doi: 10.1016/j.jviromet.2013.10.026.

31. Ren, H., Ferguson, B. J., Maluquer de Motes, C., Sumner, R. P., Harman, L. E., and Smith, G. L. Enhancement of CD8(+) T-cell memory by removal of a vaccinia virus nuclear factor-κB inhibitor. Immunology (2015) 145(1):34–49. doi: 10.1111/imm.12422.

32. Lynch, H. E., Goldberg, G. L., Chidgey, A., Van den Brink, M. R., Boyd, R., and Sempowski, G. D. Thymic involution and immune reconstitution. Trends Immunol (2009) 30(7):366–373. doi: 10.1016/j.it.2009.04.003.

33. Youm, Y. H., Horvath, T. L., Mangelsdorf, D. J., Kliewer, S. A., and Dixit, V. D. Prolongevity hormone FGF21 protects against immune senescence by delaying age-related thymic involution. Proc Natl Acad Sci U S A (2016) 113(4):1026–1031. doi: 10.1073/pnas.1514511113.

34. Pütz, M. M., Alberini, I., Midgley, C. M., Manini, I., Montomoli, E., and Smith, G. L. Prevalence of antibodies to Vaccinia virus after smallpox vaccination in Italy. J Gen Virol (2005) 86(Pt 11):2955–2960. doi: 10.1099/vir.0.81265-0.

35. Ben-Yehuda, A., Ehleiter, D., Hu, A. R., and Weksler, M. E. Recombinant vaccinia virus expressing the PR/8 influenza hemagglutinin gene overcomes the impaired immune response and increased susceptibility of old mice to influenza infection. J Infect Dis (1993) 168(2):352–357.

36. Albarnaz, J. D., Torres, A. A., and Smith, G. L. Modulating Vaccinia Virus Immunomodulators to Improve Immunological Memory. Viruses (2018) 10(310.3390/v10030101.

37. de Freitas, L. F. D., Oliveira, R. P., Miranda, M. C. G., Rocha, R. P., Barbosa-Stancioli, E. F., Faria, A. M. C. et al. The Virulence of Different Vaccinia Virus Strains Is Directly Proportional to Their Ability To Downmodulate Specific Cell-Mediated Immune Compartments. J Virol (2019) 93(610.1128/JVI.02191-18.

38. Ramírez, J. C., Gherardi, M. M., and Esteban, M. Biology of attenuated modified vaccinia virus Ankara recombinant vector in mice: virus fate and activation of B- and T-cell immune responses in comparison with the Western Reserve strain and advantages as a vaccine. J Virol (2000) 74(2):923–933.

39. Frasca, D., and Blomberg, B. B. Inflammaging decreases adaptive and innate immune responses in mice and humans. Biogerontology (2016) 17(1):7–19. doi: 10.1007/s10522-015-9578-8.

40. Bektas, A., Schurman, S. H., Sen, R., and Ferrucci, L. Human T cell immunosenescence and inflammation in aging. J Leukoc Biol (2017) 102(4):977–988. doi: 10.1189/jlb.3RI0716-335R.

41. Tilstra, J. S., Clauson, C. L., Niedernhofer, L. J., and Robbins, P. D. NF-κB in Aging and Disease. Aging Dis (2011) 2(6):449–465.

42. Minciullo, P. L., Catalano, A., Mandraffino, G., Casciaro, M., Crucitti, A., Maltese, G. et al. Inflammaging and Anti-Inflammaging: The Role of Cytokines in Extreme Longevity. Arch Immunol Ther Exp (Warsz) (2016) 64(2):111–126. doi: 10.1007/s00005-015-0377-3.

43. Franceschi, C., Garagnani, P., Vitale, G., Capri, M., and Salvioli, S. Inflammaging and ‘Garb-aging’. Trends Endocrinol Metab (2017) 28(3):199–212. doi: 10.1016/j.tem.2016.09.005.

44. Franceschi, C., Garagnani, P., Parini, P., Giuliani, C., and Santoro, A. Inflammaging: a new immune-metabolic viewpoint for age-related diseases. Nat Rev Endocrinol (2018) 14(10):576–590. doi: 10.1038/s41574-018-0059-4.

45. Gupta, S., Termini, J. M., Kanagavelu, S., and Stone, G. W. Design of vaccine adjuvants incorporating TNF superfamily ligands and TNF superfamily molecular mimics. Immunol Res (2013) 57(1-3):303–310. doi: 10.1007/s12026-013-8443-6.

46. Luo, J., Zhang, B., Wu, Y., Tian, Q., Zhao, J., Lyu, Z. et al. Expression of interleukin-6 by a recombinant rabies virus enhances its immunogenicity as a potential vaccine. Vaccine (2017) 35(6):938–944. doi: 10.1016/j.vaccine.2016.12.069.

47. Chen, G., Lustig, A., and Weng, N. P. T cell aging: a review of the transcriptional changes determined from genome-wide analysis. Front Immunol (2013) 4121. doi: 10.3389/fimmu.2013.00121.

48. Vescovini, R., Biasini, C., Fagnoni, F. F., Telera, A. R., Zanlari, L., Pedrazzoni, M. et al. Massive load of functional effector CD4+ and CD8+ T cells against cytomegalovirus in very old subjects. J Immunol (2007) 179(6):4283–4291.

49. Weng, N. P., Akbar, A. N., and Goronzy, J. CD28(-) T cells: their role in the age-associated decline of immune function. Trends Immunol (2009) 30(7):306–312. doi: 10.1016/j.it.2009.03.013.

50. Franceschi, C., Salvioli, S., Garagnani, P., de Eguileor, M., Monti, D., and Capri, M. Immunobiography and the Heterogeneity of Immune Responses in the Elderly: A Focus on Inflammaging and Trained Immunity. Front Immunol (2017) 8982. doi: 10.3389/fimmu.2017.00982.

51. Fulop, T., Larbi, A., Dupuis, G., Le Page, A., Frost, E. H., Cohen, A. A. et al. Immunosenescence and Inflamm-Aging As Two Sides of the Same Coin: Friends or Foes. Front Immunol (2017) 81960. doi: 10.3389/fimmu.2017.01960.

