## Supplementary Data for "Enhanced efficacy of vaccination with vaccinia virus in old versus young mice"

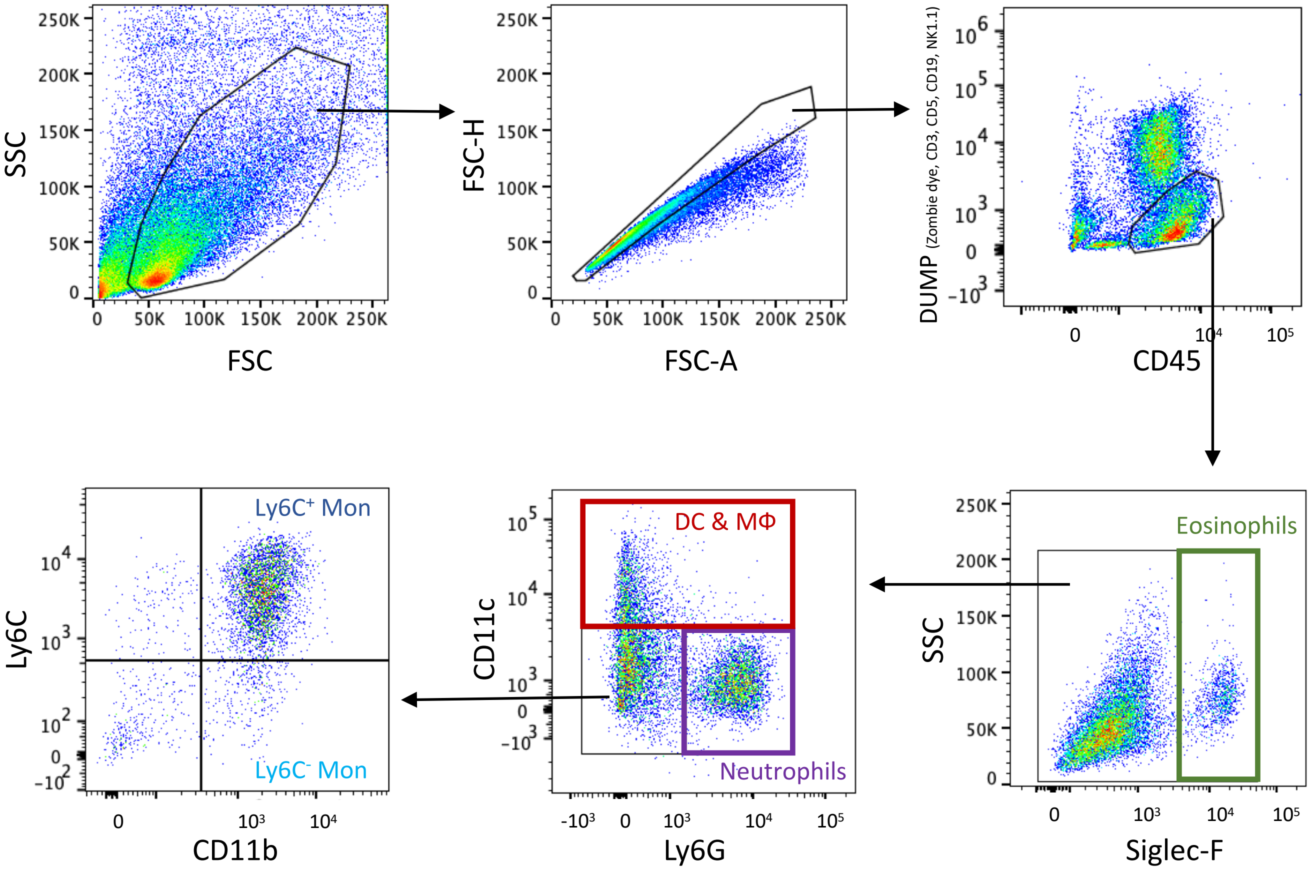


**Supplementary figure 1**. Gating strategy for flow cytometry of myeloid lineage cells present in mouse ear tissue 7 d after intradermal injection with $10^4$ PFU of VACV strain WR. Cells were gated on their ability to scatter light. Doublets were excluded using a FSC-A versus FSC-H plot. The myeloid gate included CD45$^{+}$Zombie dye$^{-}$CD3$^{-}$CD5$^{-}$CD19$^{-}$NK1.1$^{-}$ cells. Further myeloid cell subpopulations were classified as follows: Eosinophils: CD45$^{+}$CD3$^{-}$CD5$^{-}$CD19$^{-}$NK1.1$^{-}$CD11c$^{-}$ Siglec- F$^{+}$; DC and MΦ (dendritic cells and macrophages): CD45$^{+}$CD3$^{-}$CD5$^{-}$CD19$^{-}$NK1.1$^{-}$Siglec-F$^{-}$ CD11c$^{+}$; Neutrophils: CD45$^{+}$CD3$^{-}$CD5$^{-}$CD19$^{-}$NK1.1$^{-}$CD11c$^{-}$Siglec-F$^{-}$ Ly6G$^{+}$; Ly6C$^{+}$Mon (inflammatory monocytes): CD45$^{+}$CD3$^{-}$CD5$^{-}$CD19$^{-}$NK1.1$^{-}$CD11c$^{-}$Siglec-F$^{-}$Ly6G$^{-}$CD11b$^{+}$Ly6C$^{+}$; Ly6C$^{-}$Mon (residential monocytes): CD45$^{+}$CD3$^{-}$CD5$^{-}$CD19$^{-}$NK1.1$^{-}$CD11c$^{-}$Siglec-F$^{-}$Ly6G$^{-}$ CD11b$^{+}$Ly6C$^{-}$.

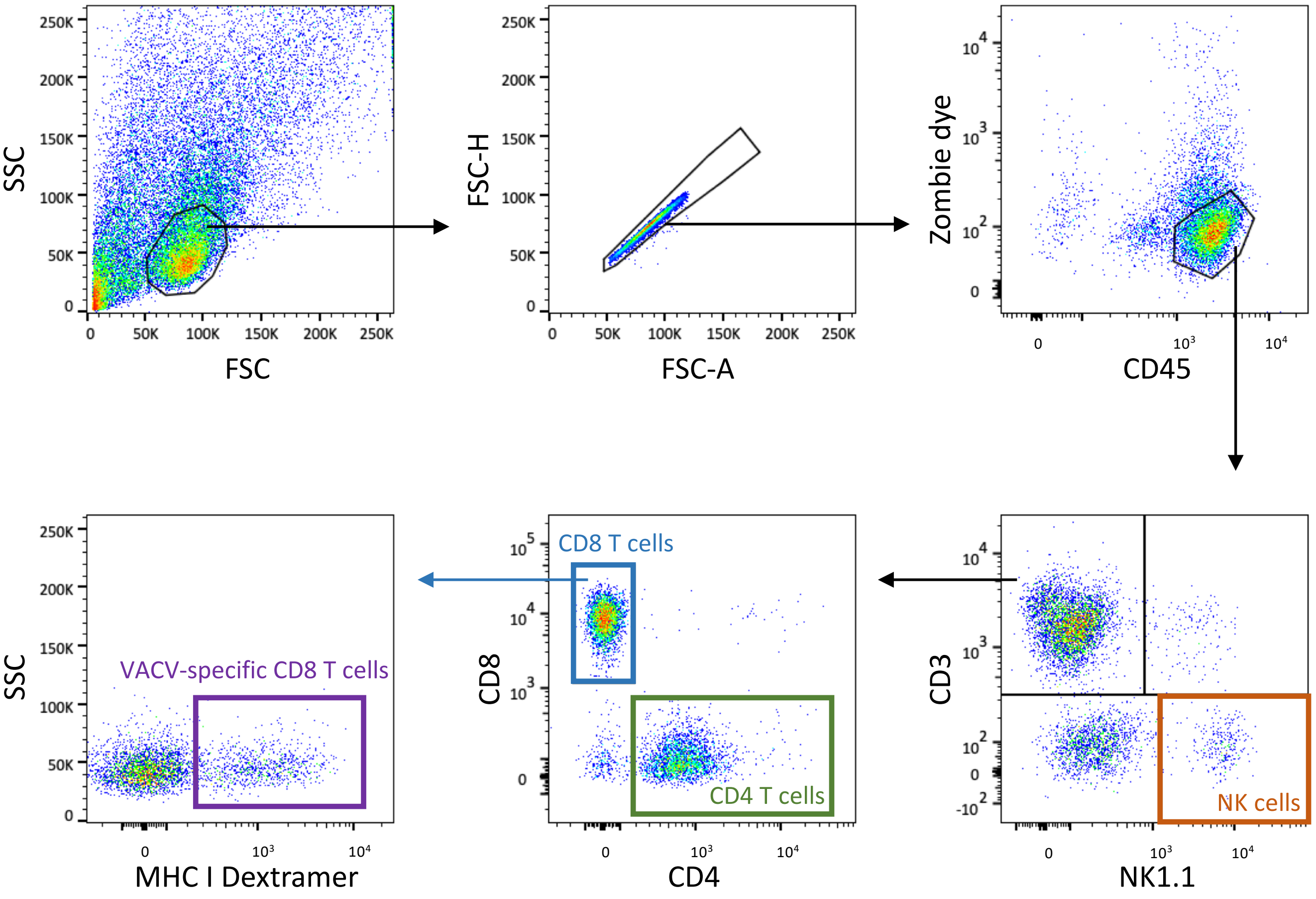


**Supplementary figure 2**. Gating strategy for flow cytometry of lymphoid lineage cells present in mouse ear tissue 7 d after intradermal injection with $10^4$ PFU of VACV strain WR. Cells were gated on their ability to scatter light. Doublets were excluded using a FSC-A versus FSC-H plot. Then, hemopoietic cells were gated as $CD45^+$Zombie dye$^-$ cells. Further lymphoid subpopulations were classified as follows: NK cells: $CD45^+CD3^-NK1.1^+$; CD4 T cells: $CD45^+CD3^+NK1.1^-CD8^-CD4^+$; CD8 T cells: $CD45^+CD3^+NK1.1^-CD4^-CD8^+$; VACV-specific CD8 T cells: $CD45^+CD3^+NK1.1^-CD4^-CD8^+$Dextramer$^+$.

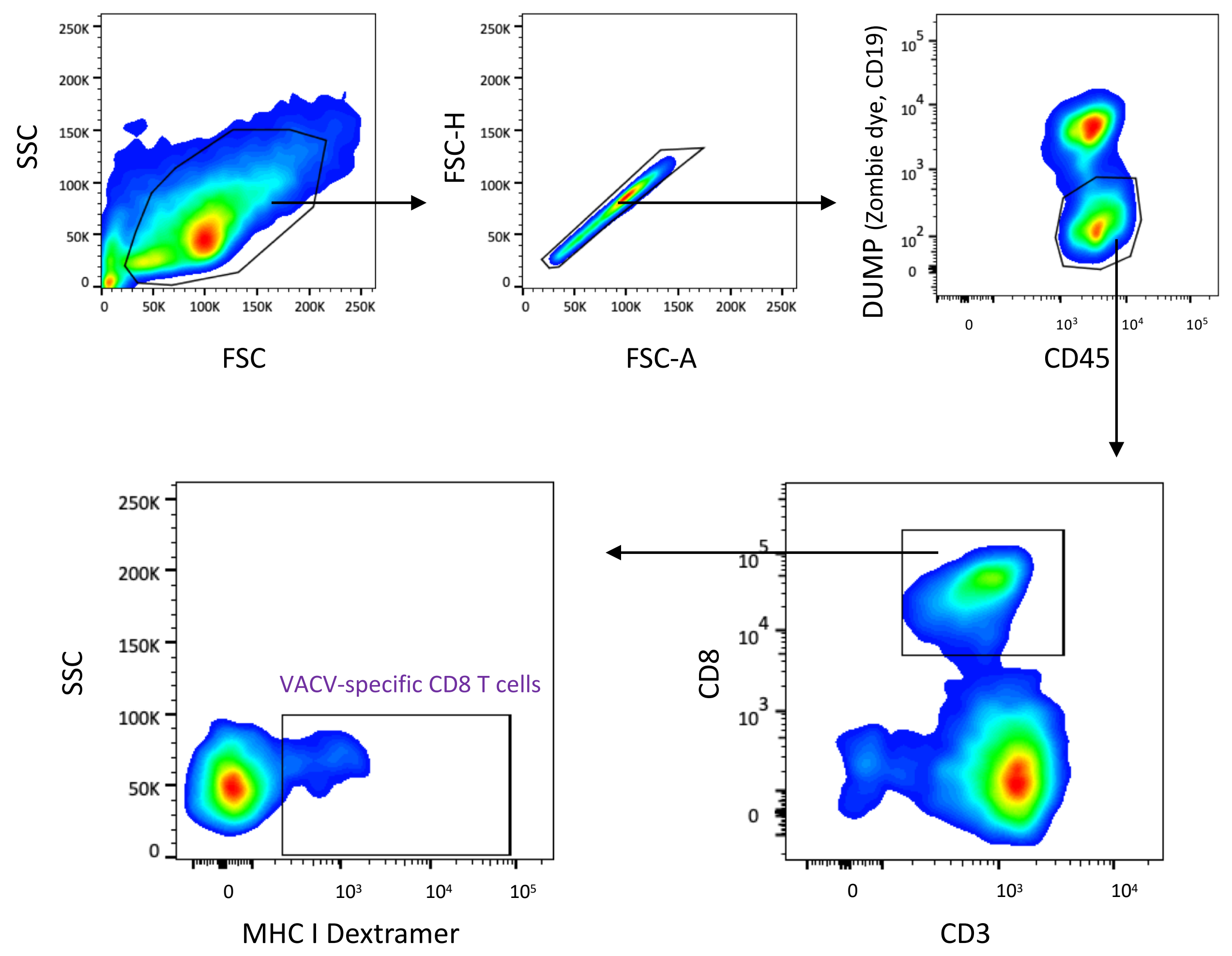


**Supplementary figure 3**. Gating strategy for flow cytometry of VACV-specific CD8 T cells obtained from draining cervical lymph nodes 7 d post intradermal injection of mouse ear pinnae with $10^4$ PFU of VACV strain WR. Cells were gated on their ability to scatter light, and doublets were excluded using a FSC-A versus FSC-H plot. Then, VACV-specific CD8 T lymphocytes were gated as $CD45^+CD3^+NK1.1^-CD4^-CD8^+Dextramer^+$cells.

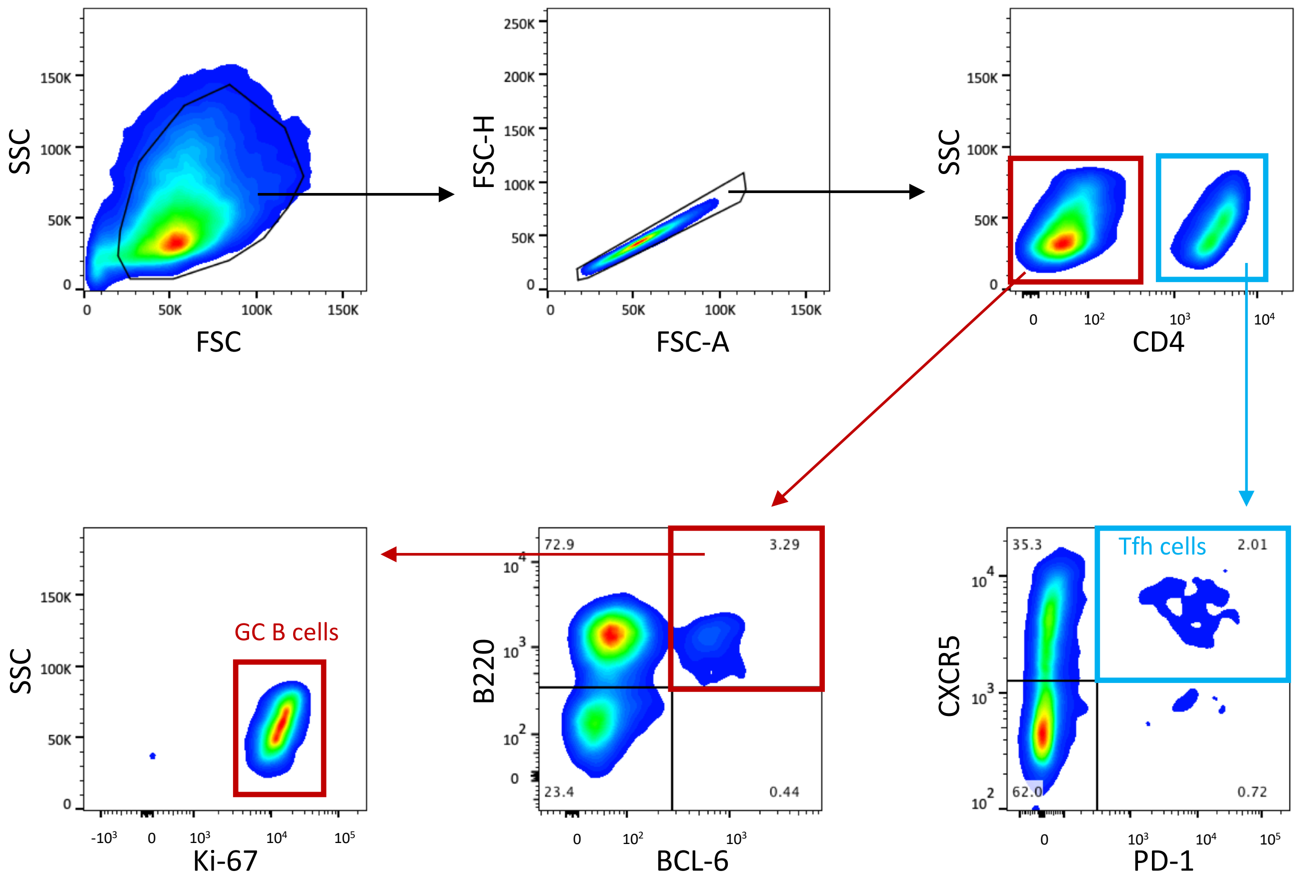


**Supplementary figure 4**. Gating strategy for flow cytometry of germinal centre (GC) B cells and T follicular helper (Tfh) lymphocytes from draining cervical lymph nodes 7 d post intradermal infection of mouse ear pinnae with $10^4$ PFU of VACV strain WR. Cells were gated on their ability to scatter light. Doublets were excluded using a FSC-A versus FSC-H plot. Further lymphoid subpopulations were classified as follows: GC B cells: $CD4^-B220^+BCL\text{-}6^+Ki\text{-}67^+$; Tfh cells: $CD4^+CXCR5^+PD\text{-}1^+$.

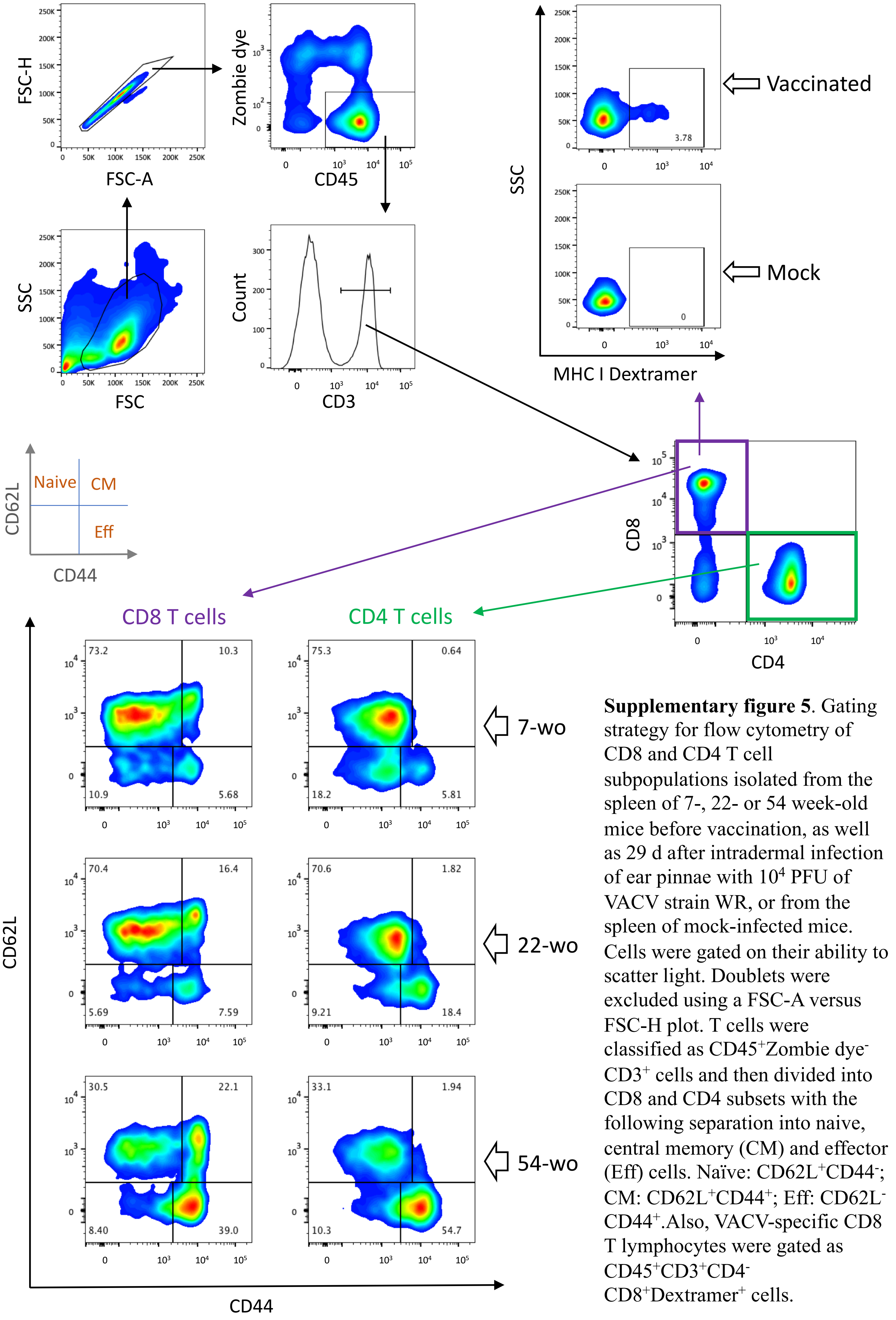


**Supplementary figure 5**. Gating strategy for flow cytometry of CD8 and CD4 T cell subpopulations isolated from the spleen of 7-, 22- or 54 week-old mice before vaccination, as well as 29 d after intradermal infection of ear pinnae with $10^4$ PFU of VACV strain WR, or from the spleen of mock-infected mice. Cells were gated on their ability to scatter light. Doublets were excluded using a FSC-A versus FSC-H plot. T cells were classified as $CD45^+$Zombie $dye^-$ $CD3^+$ cells and then divided into CD8 and CD4 subsets with the following separation into naive, central memory (CM) and effector (Eff) cells. Naïve: $CD62L^+CD44^-$; CM: $CD62L^+CD44^+$; Eff: $CD62L^-$ $CD44^+$.Also, VACV-specific CD8 T lymphocytes were gated as $CD45^+CD3^+CD4^-$ $CD8^+Dextramer^+$ cells.

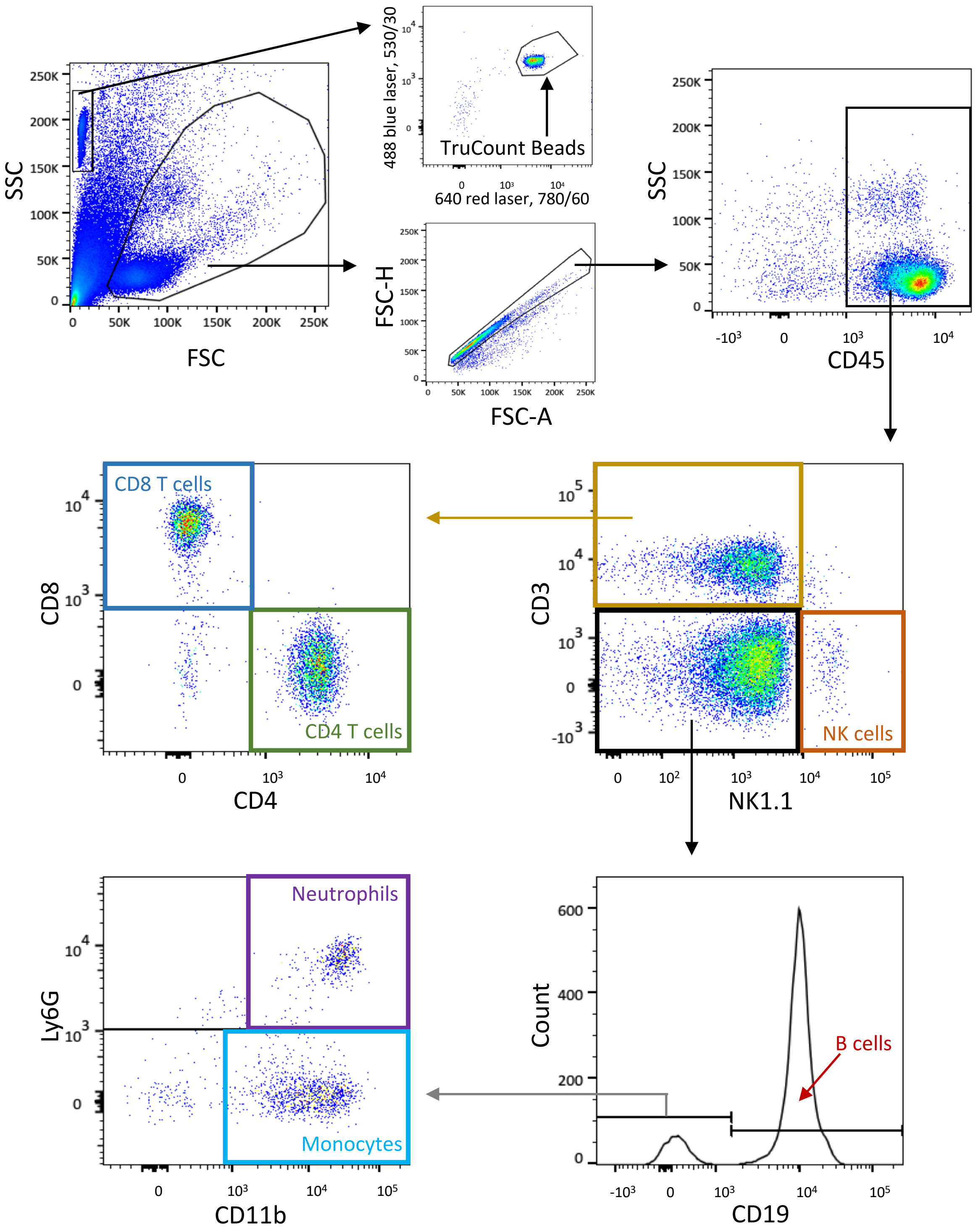


**Supplementary figure 6**. Gating strategy for Trucount flow cytometry of different leukocyte populations present in mouse blood. First, cell and bead areas were identified based on their ability to scatter light. Then the beads were gated based on their bright fluorescence in a variety of channels. Cell doublets were excluded using a FSC-A versus FSC-H plot, then hemopoietic cells were gated as CD45+cells. Further leukocyte subpopulations were classified as follows: NK: $CD45^+CD3^-NK1.1^+$; CD4 T cells: $CD45^+CD3^+NK1.1^-CD8^-CD4^+$; CD8 T cells: $CD45^+CD3^+NK1.1^-CD4^-CD8^+$; B cells: $CD45^+CD3^-NK1.1^-CD19^+$; Neutrophils: $CD45^+CD3^-NK1.1^-CD19^-CD11b^+Ly6G^+$; Monocytes: $CD45^+CD3^-NK1.1^-CD19^-CD11b^+Ly6G^-$.

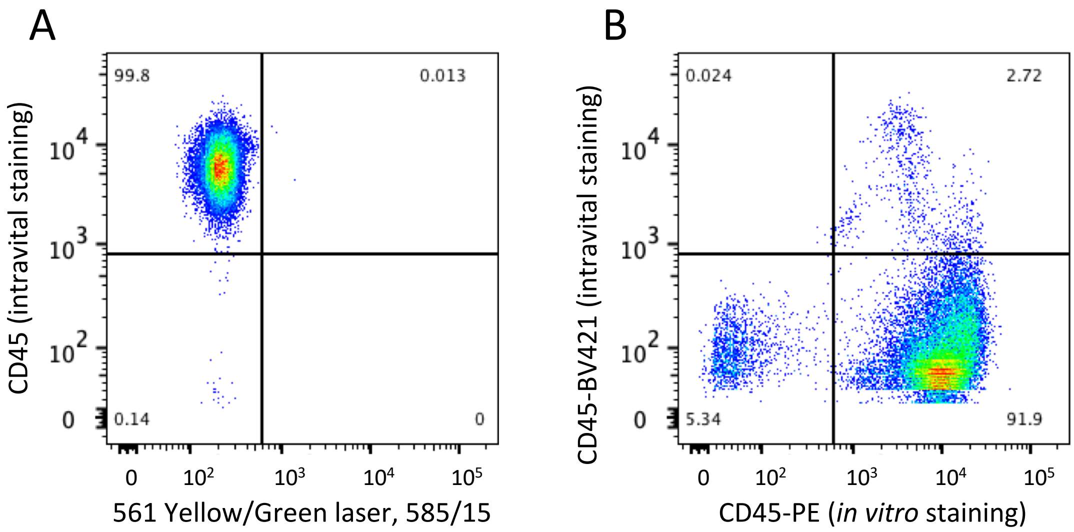


**Supplementary figure 7.** Intravascular and *in vitro* staining with anti-CD45 antibodies. (**A**) Flow cytometry of blood sample after intravascular staining with anti-CD45 antibodies. (**B**) Flow cytometry of cells from the ear of a mouse 7 d after intradermal infection with $10^4$ PFU of VACV strain WR after intravascular staining with anti-CD45-BV421 antibodies followed by *in vitro* staining with anti-CD45-PE antibodies.

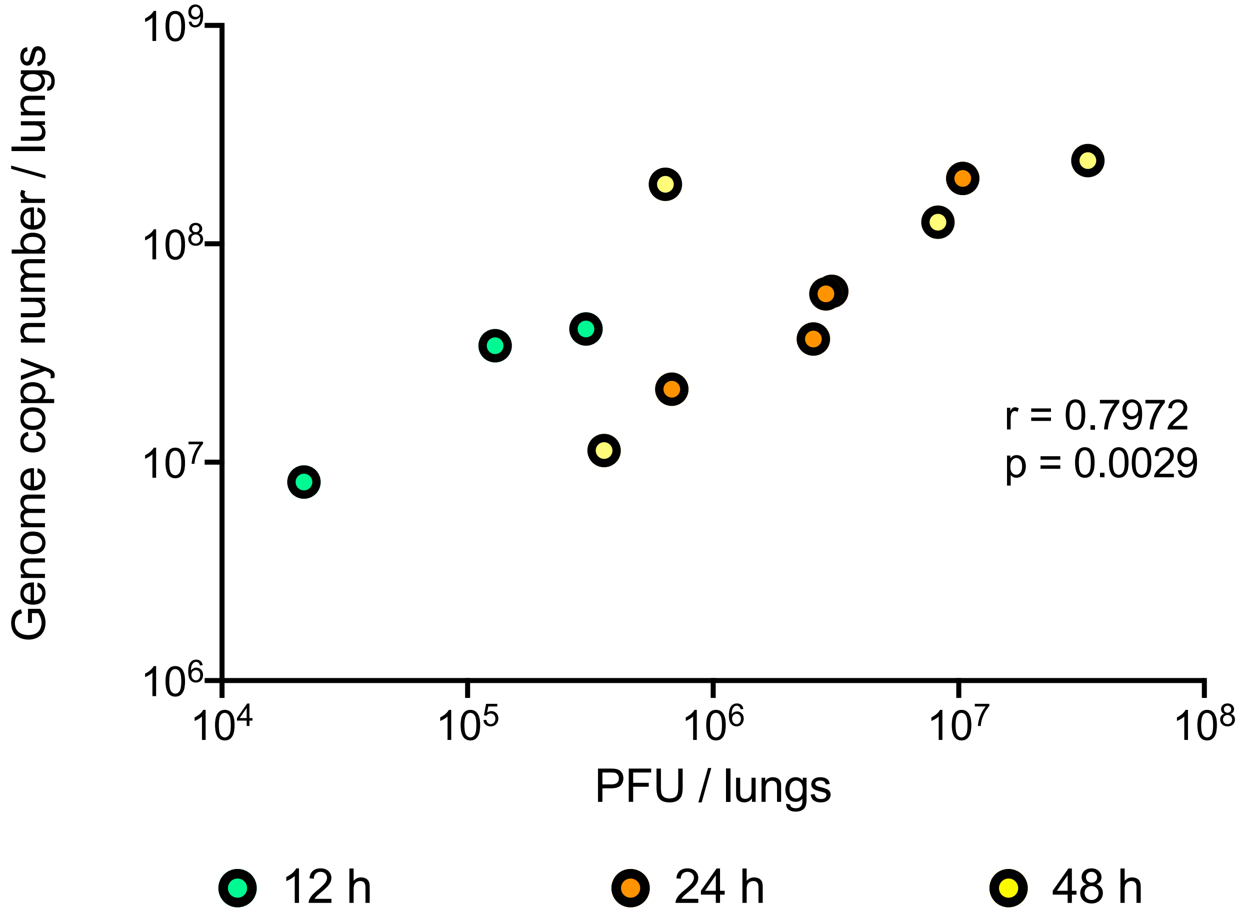


**Supplementary figure 8.** Spearman correlation analysis of infectious virus titer versus virus genome copy number. VACV loads were measured in the lungs of naïve mice at 12, 24, or 48 h after intranasal infection with $0.7 \times 10^7$ PFU of VACV WR by either plaque assay on BSC-1 cells (expressed as plaque-forming units [PFU]/ lungs), or by qPCR using primers for the VACV *E9L* gene (expressed as genome copy number / lungs).

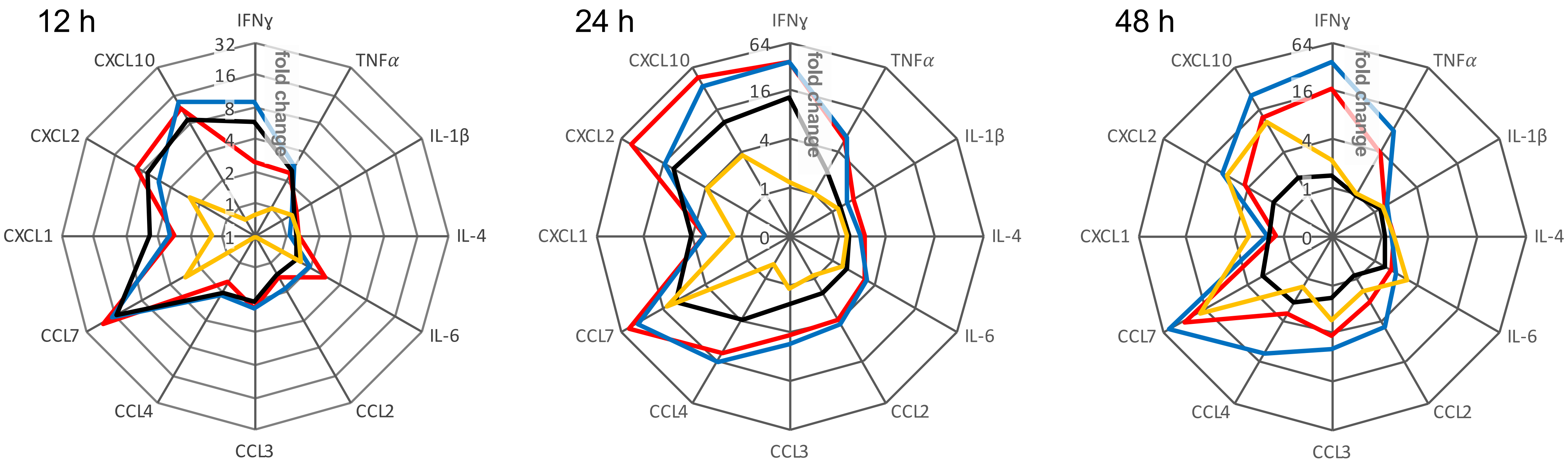


**Supplementary figure 9.** Alternative representation of data from Fig. 7. Groups (*n*=4-5) of 7-, 22- or 54-wo C57BL/6 mice were infected intradermally with $10^4$ PFU of VACV WR. After 33 d, the mice were infected intranasally with VACV WR and the lungs were collected at 12, 24 and 48 h later. The levels of cytokines and chemokines were measured by multiplex assay (Luminex). The radar plots display data from the three age groups (7-wo in red, 22-wo in blue and 54-wo in black) together with naive (mock-vaccinated) animals (in yellow). Data presented are the fold change from the baseline levels of each cytokine/chemokine in mice before intranasal challenge. Means are shown.

**Supplementary table 1.** Monoclonal antibodies and dyes used for staining cells prior to analysis by flow cytometry.

| Antibody/dye | Clone | Source |
|---|---|---|
| B220-PerCP | RA3-6B2 | 103234, BioLegend |
| Bcl-6-PE | K112-91 | 561522, BD Biosciences |
| CD3-APC | 145-2C11 | 553066, BD Biosciences |
| CD3-BV421 | 145-2C11 | 562600, BD Biosciences |
| CD4-APC-H7 | GK1.5 | 560181, BD Biosciences |
| CD4-BUV395 | GK1.5 | 563790, BD Biosciences |
| CD5-BV421 | 53-7.3 | 562739, BD Biosciences |
| CD8-BB515 | 53-6.7 | 564422, BD Biosciences |
| CD8-BV605 | 53-6.7 | 100744, BioLegend |
| CD11b-PE | M1/70 | 101208, BioLegend |
| CD11c-BV650 | N418 | 117339, BioLegend |
| CD19-BV421 | 1D3 | 562701, BD Biosciences |
| CD44-BB515 | IM7 | 564587, BD Biosciences |
| CD45-PerCP | 30-F11 | 557235, BD Biosciences |
| CD62L-APC-C7 | MEL-14 | 104428, BioLegend |
| CXCR5-BV421 | L138D7 | 145512, BioLegend |
| Ki-67-FITC | SolA15 | 11-5698-82, eBioscience |
| Ly6C-APC | HK1.4 | 128016, BioLegend |
| Ly6G-APC-H7 | 1A8 | 565369, BD Biosciences |
| MHC Dextramer H-2Kb/TSYKFESV/PE | | JD3267-PE, Immudex |
| NK1.1-BV421 | PK136 | 562921, BD Biosciences |
| NK1.1-BV605 | PK136 | 108739, BioLegend |
| PD-1-APC (J43) | J43 | 562671, BD Biosciences |
| Siglec-F-BB515 | E50-2440 | 564514, BD Biosciences |
| Zombie Green Fixable Viability Kit | | 423111, BioLegend |
| Zombie Violet Fixable Viability Kit | | 423113, BioLegend |
| PE Mouse IgG1, κ Isotype Control | MOPC-21 | 554680, BD Biosciences |
| FITC Rat IgG2a kappa Isotype Control | eBR2a | 11-4321-80, eBioscience |
